# Genomic Convergence in Hibernating Mammals Elucidates the Genetics of Metabolic Regulation in the Hypothalamus

**DOI:** 10.1101/2024.06.26.600891

**Authors:** Elliott Ferris, Josue D. Gonzalez Murcia, Adriana Cristina Rodriguez, Susan Steinwand, Cornelia Stacher Hörndli, Dimitri Traenkner, Pablo J Maldonado-Catala, Christopher Gregg

**Author notes:** These authors contributed equally to this work (Josue D. Gonzalez Murcia, Adriana Cristina Rodriguez).

## Abstract

Elucidating the genetic basis of mammalian metabolism could help define mechanisms central to health and disease. Here, we define conserved *cis-*regulatory elements (CREs) and programs for mammalian metabolic control. We delineate gene expression and chromatin responses in the mouse hypothalamus for 7 steps of the Fed-to-Fasted-to-Refed (FFR) response process. Comparative genomics of hibernating versus non-hibernating lineages then illuminates *cis-*elements showing convergent changes in hibernators. Hibernators accumulated loss-of-function effects for specific CREs regulating hypothalamic FFR responses. Multi-omics approaches pinpoint key CREs, genes, regulatory programs, and cell types in the divergence of hibernating and homeothermic lineages. The refeeding period after extended fasting is revealed as one critical period of chromatin remodeling with convergent genomic changes. This genetic framework is a step toward harnessing hibernator adaptations in medicine.

**One sentence summary:** Convergent signals define *cis-*regulatory mechanisms behind food scarcity responses and hibernator-homeotherm divergence.

---

Metabolism encompasses the essential chemical processes for cell and organism function and its proper regulation is crucial. Metabolic imbalances contribute to diseases such as neurodegenerative disease (*1*), obesity (*2*, *3*), and cancer (*4*). On the other hand, interventions influencing metabolism can aid in disease prevention and recovery, and promote longevity (*5*, *6*). Species differences in metabolic traits could be leveraged to uncover the genetic control of mammalian metabolism. Hibernation constitutes an extreme metabolic phenotype (*7*, *8*). Obligate hibernators evolved a seasonal regulation of behavior and metabolism in which their body mass increases by 30%–50% (*9*) and they enter long periods of torpor sustained by stored fat during seasons of food scarcity (*7*, *8*). On the other hand, facultative hibernators respond to acute food deprivation or cold stresses with short periods of torpor with suppressed metabolism and body temperature (*7*, *10*), while homeothermic mammals are incapable of such dramatic metabolic changes and maintain more stable metabolic rates and body temperatures through different seasons and environmental conditions (*7*, *8*). Studies of different hibernator adaptations suggest that changing components of mammalian metabolism offers benefits such as metabolic rate and feeding control, neuroprotection (*11*), reversal of neurodegenerative processes (*12–14*), obesity and insulin resistance control (*9*, *15*, *16*), tumor dormancy (*17*, *18*), and enhanced longevity (*19*). Uncovering the genetic mechanisms involved is expected to provide important advances in our understanding of mammalian metabolism and health.

Hibernation has arisen independently in different species across at least 7 orders of mammals, ranging from monotremes to primates (*20*). The repeated appearance of hibernation in different lineages suggests that most of the protein-coding genes involved are present in both hibernators and non-hibernators, with gene regulatory modifications likely accounting the profound phenotypic differences. Changes to *cis-*regulatory elements (CREs) are primary genetic determinants of phenotypic diversity in evolution (*21*, *22*), as well as common human disease (*23*). Computationally solving the *cis-*regulatory programs underlying complex traits has proven difficult because the genome lacks the sequence diversity needed for models to learn the relevant parameters (*24*), indicating that targeted, experimental approaches are vital. We previously showed that conserved genetic elements exhibiting accelerated evolution in lineages with biomedically important traits point to candidate *cis-*regulatory mechanisms for those traits, including convergent genomic effects on obesity risk loci in hibernators (*25*, *26*). These, and other studies (*27–29*), indicate that convergent evolutionary changes in independent lineages with related phenotypes are effective for uncovering genetic mechanisms for complex traits.

Here, we use comparative genomics approaches to uncover *cis-*regulatory elements (CREs) regulating mammalian metabolism and hibernation-relevant phenotypes. Our approach concentrates on the hypothalamus because it is a central hub of metabolic and physiological control (*30*). The hypothalamus harbors a myriad of specialized cell types that govern metabolism, feeding, activity, thermogenesis, energy expenditure, torpor, and more (*30–32*), yet the *cis-*regulatory programs involved are largely undefined. Utilizing genetically tractable murine models (*33*) capable of brief fasting-induced torpor bouts and changes to metabolic rate (*33–36*), we assess gene and chromatin dynamics across various fed, fasted, and refed (FFR) states to define hypothalamic genetic programs for FFR responses. Leveraging a genome alignment from the Zoonomia consortium (*37*), we then define regions of the mammalian genome conserved in homeotherms but show convergent accelerated evolution in lineages that independently evolved hibernation. Convergent genomic changes in hibernators affect *cis-*elements that contact hub genes central to FFR responses. We characterize these elements at the cellular level, uncovering key CREs, hypothalamic cell-types, and gene regulatory programs. Our companion study shows functional roles for individual CREs identified with this approach in modulating metabolism and behavior (*38*). Our studies unveil *cis-*regulatory mechanisms underlying mammalian metabolic control.

## Genetic Programs for Fed-Fasted-Refed (FFR) Responses in the Mouse Hypothalamus

Hibernation is an adaptation for surviving periods of food scarcity (**Fig. 1A**). We hypothesized that hibernator lineages evolved distinct hypothalamic genetic programs impacting FFR responses (**Fig. 1A**). We first characterized metabolic and behavioral responses to different FFR states in mice, which show bouts of fasting-inducible torpor (*33*). Adult female mice were implanted with small internal body temperature (Tb_i_) monitors and placed in CLAMS (comprehensive lab animal monitoring system) metabolic cages to measure changes during a 10-day FFR paradigm that includes 48hrs in a fed baseline state (Fed phase), 48hrs of food deprivation (FD) + reduced ambient temperature (shift from 25°C to 18°C) for torpor induction (Torpor phase), and 6 days of refeeding (RF) at 25°C with ad libitum food (RF phase) (**Fig. 1B and S1A**). At baseline, mice have a Tb_i_ of 38°C and 36°C during the dark versus light cycles, respectively. The Tb_i_ drops in response to FD + reduced ambient temperature, indicative of torpor bouts, reaching as low as ∼20°C. Within hours of RF, Tb_i_ in the Fed and RF phases were indistinguishable. The Respiratory Exchange Ratio (RER) significantly increased during the post-torpor RF phase and remained elevated for at least 6 days, reaching levels that are >1, which indicates excessive CO_2_ production and energy consumption (**Fig. 1B**). Metabolic rate, locomotor activity, and energy expenditure are significantly decreased during RF compared to the Fed period **(Fig. S1A-C)**, and the decrease in metabolic rate and locomotion persists for at least 6 days **(Fig. S1A,B).** We learned that the RF phase is an extended period of metabolic and behavioral change after torpor. We then set out to define hypothalamic genetic programs to learn how they changed in hibernators.

**Fig. 1.**
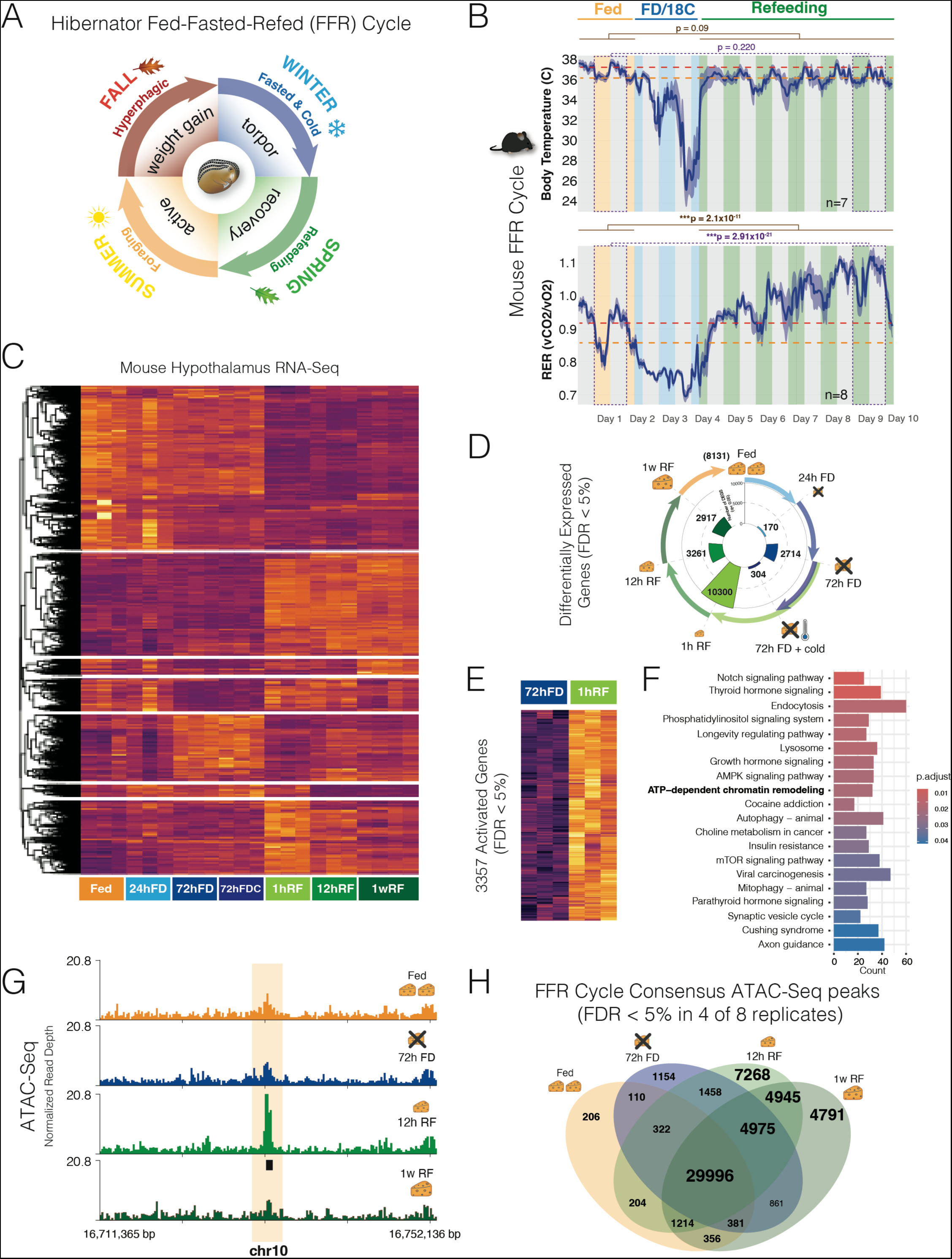
Gene expression programs and chromatin accessibility dynamics uncovered for discrete stages of a Fed-Fasted-Refed (FFR) Response in the mouse hypothalamus. **(A)** Schematic summary of the Fed-Fasted-Refed (FFR) cycle experienced by obligate hibernators, like the 13-lined ground squirrel. Different metabolic states and processes are induced during the FFR cycle. Facultative hibernators show torpor responses to food deprivation and/or cold, rather than seasonal cycles. **(B)** Internal body temperature and respiratory exchange ratio (RER) in adult female mice during Fed (ambient temperature of 24°C) (yellow), food deprived (FD) + cold (ambient temperature of 18°C) (blue), and Refed states (green) over 10 days in CLAMS cages. RER significantly differs for Fed and Refed states (generalized linear model, brown), with increased RER during refeeding, including for the day 1 versus day 9 comparison (purple). Red dashed line, mean for dark cycle (grey bars). Orange dashed line, mean for light cycle (colored bars). Dark blue line and shading, mean±SEM, n=10. Also see Fig. S1. **(C)** Heatmap of RNA-Seq measured gene expression levels in the adult mouse hypothalamus for genes differentially expressed according to FFR state (FDR <5%, n=3-4). Shows counts per million scaled by column. FD, food deprived; RF, refed; 12hrs RF, refed for 12hrs after 72hrs of FD; 1w RF, refed for 1 week after 72hrs of FD; h, hours; w, week; Cold is ambient exposure to 18°C compared to 24°C. **(D)** Circular plot shows the numbers of significant differentially expressed genes at progressive steps of the FFR cycle detected by RNA-Seq in mouse hypothalamus of the mouse. Colors of the bars and arrows match and indicate gene expression comparisons for each step in the cycle. Numbers of significant DEGs (differentially expressed gene, FDR < 5%) are shown in the inner circle. **(E and F)** The heatmap shows DEGs that significantly increase expression from 72hr FD to 1hr RF (E) and the barplot shows the KEGG pathways significantly enriched for these RF increased DEGs (FDR < 5%) (F). **(G)** The signal plots show an example of an intergenic ATAC-Seq consensus peak (FDR < 5% in 4 of 8 samples) identified in the 12hr RF period and not the Fed, 72hr FD or 1 week RF stages in the adult mouse hypothalamus (n=8). **(H)** Venn diagram depicting numbers of consensus ATAC-Seq peaks genome-wide in the hypothalamus, highlighting a rise in chromatin accessibility sites in refeeding (RF) conditions versus Fed and 72-hour fasted (FD) states. Consensus peaks are significant in at least 4 of 8 biological replicates (FDR < 5%, n=8).

To examine gene expression programs, we performed RNA-Seq profiling of the hypothalamus for 7 comparisons across the FFR response. These comparisons included: (**1**) fed baseline versus 24hr FD; (**2**) 24hr FD versus 72hr FD; (**3**) 72hr FD versus 72hr FD + cold (18°C); (**4**) 72hr FD versus 1hr RF; (**5**) 1hr RF versus 12hrs RF; (**6**) 12hrs RF versus 1 week RF; and (**7**) 1 week RF versus Fed baseline. Modest gene expression changes occurred for the step from Fed to 24hr FD, while the step from 24hr FD to 72hr FD affected thousands of genes and combining a reduced ambient temperate with 72hr FD affected 304 additional genes (**Fig. 1C and D**). RF significantly affected the expression of over ten thousand genes (FDR < 5%) within 1hr of RF relative to 72hr FD (**Fig. 1D**). The step from 1hr to 12hrs of RF significantly affected 3261 genes and the step to 1 week of RF affected 2917 genes. Finally, as expected from our CLAMS data, mice did not return to a Fed baseline state after 1 week and thousands of genes showed significant differences in expression in 1 week RF versus Fed baseline mice (**Fig. 1D**). Thus, we found that extended FD (72hr FD) is required to induce substantial hypothalamic gene expression changes, and that the RF period involves various gene expression changes over many days (**Table S1**).

A comparative biological process, reactome pathway, KEGG pathway, and disease ontology gene set enrichment analysis of the FFR response DEGs uncovered biological processes, pathways, and disease mechanisms significantly activated or suppressed at each step of the FFR program (**Fig. S2A and Table S2**). The analyses yielded several findings. Fasting and hibernation affect neurodegeneration (*14*, *39*) and we found that neurodegeneration pathways are disproportionately suppressed during the transition from 24hr FD to 72hr FD, activated at 1hr RF, and continue to show increased expression at 1week RF compared to Fed baseline (**Fig. S2C**). Immunity and inflammatory pathways, also affected by fasting and hibernation (*40–42*), show changes at specific steps of the FFR response. For example, in the transition from 72hr FD to 72hr FD + cold (18°C), natural killer cell mediate cytotoxicity and other infectious disease response pathways are activated, showing the effects of adding cold to FD (**Fig. S2C**). Finally, the longevity regulating pathway is significantly increased during RF at the transition from 1hr to 12hr RF (**Fig. S2C**), which is noteworthy because fasting and hibernation promote longevity (*6*, *19*, *43*). Thus, steps of the FFR response impact different signaling pathways, mitochondrial processes, apoptosis, synaptic plasticity, immunity, autophagy, and more (**Table S2**).

Chromatin remodeling pathways are also affected during RF. Refeeding 1hr after 72hr FD disproportionately activated genes involved in ATP-dependent chromatin remodeling, as well as histone deacetylases and transcription regulatory genes (**Fig. 1E and F; Table S2**). These results suggested that RF after FD is a critical period of chromatin remodeling and epigenetic reprogramming. To test this and uncover CREs affected by FFR responses, we profiled chromatin accessibility in the adult mouse hypothalamus using ATAC-Seq in: (**1**) Fed baseline, (**2**) 72hr FD, (**3**) 12hr RF, and (**4**) 1 week RF mice. We uncovered an average of 50K to 70K significant ATAC-Seq peaks in each condition (FDR 5%, n=8). High confidence consensus ATAC-Seq peak sites were defined for each condition as sites with significant accessibility (FDR <5%) in at least 4 out of 8 replicates (**Table S3**). We found distal genomic regions with significant chromatin accessibility in specific FFR states (**Fig. 1G,H**). A Venn diagram analysis showed that thousands of consensus sites are unique to the RF period, including accessibility sites for the 12hr RF and 1 week RF time points (**Fig. 1H**). Therefore, RF is a critical period of hypothalamic epigenetic and gene expression changes. This understanding of the FFR response program in the mouse established foundations for exploring how evolutionary changes in obligate hibernators impacted this program.

## Convergent Evolution in Hibernators Targets the Hypothalamic FFR Genetic Program

Our foundational hypothesis posits that hibernating lineages selected for changes to conserved *cis-*elements regulating FFR responses to affect metabolic control mechanisms in the hypothalamus. With recent Zoonomia data including new species, we contrasted accelerated evolution in hibernators and homeotherms (*26*) (*44*). We tested ∼1.3 million DNA sequences conserved for over ∼100 million years in a set of background homeotherms, and referred to as conserved regions (CRs), for accelerated evolution in each of the hibernators and in control homeotherms closest to the hibernators in the multiple genome alignment. We found significant accelerated regions (ARs) in lineages that independently evolved hibernation, including the small Madagascar hedgehog (*Echinops telfairi*), little brown bat (*Myotis lucifugus*), thirteen-lined ground squirrel (*Ictidomys tridecemlineatus*), and the fat-tailed dwarf lemur (*Cheirogaleus medius*) (**Fig. 2A,B**). We also defined significant ARs in control homeotherms that are the closest lineages to the hibernators in the multiple alignment, as well as deletions in CRs in hibernators and control homeotherms (**Fig. 2A,B**). Overall, the analysis identified ARs and deleted regions across species’ genomes, showing genetic changes linked to trait evolution in each lineage.

**Fig. 2.**
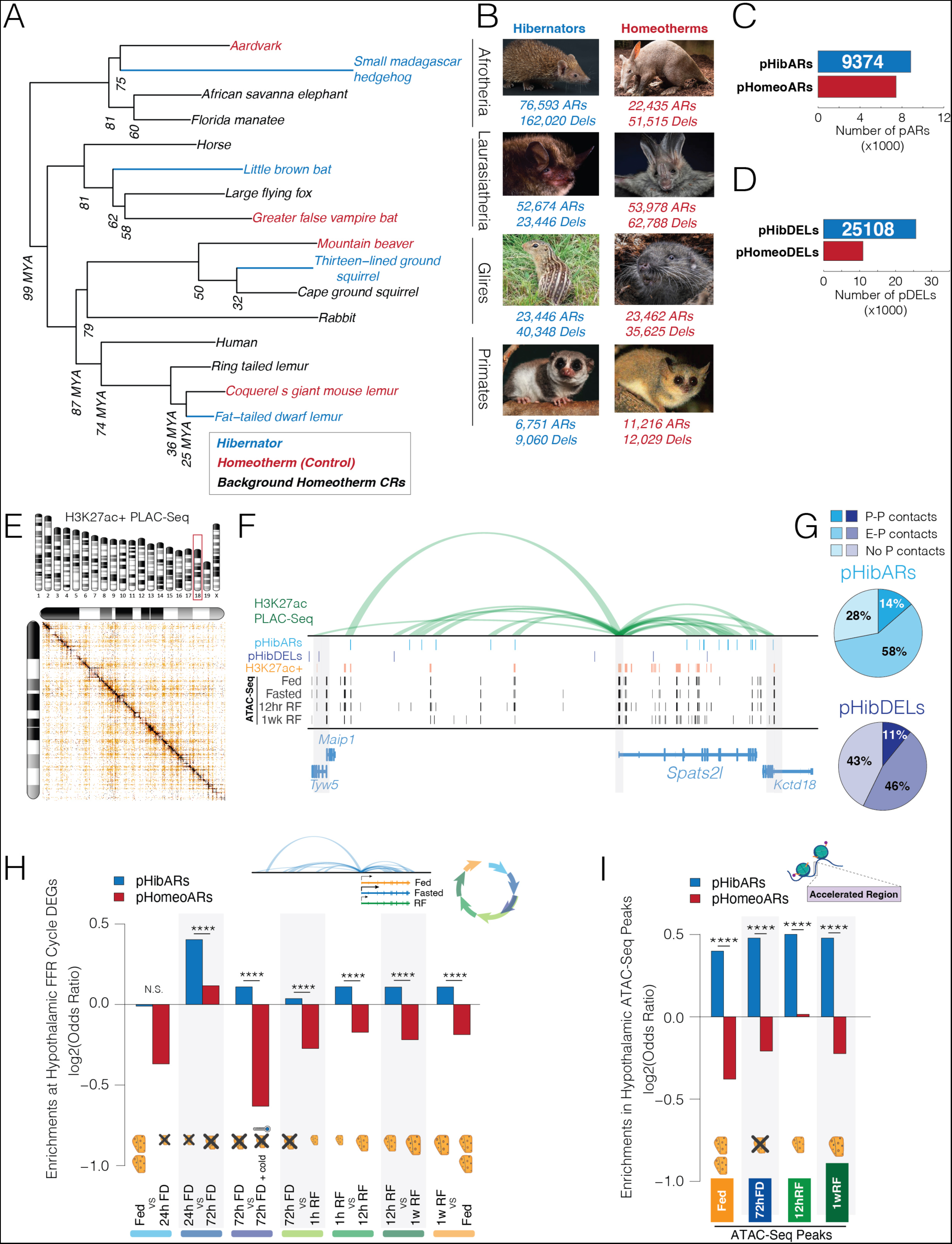
Parallel genomic changes in hibernators disproportionately impact gene regulatory mechanisms controlling hypothalamic FFR genetic programs. **(A)** Phylogenetic tree showing the lineages used in our study from the 241-way mammalian species multiple alignment (241-mammalian-2020v2b.maf). We identified obligate hibernators in different clades (blue) and homeotherm controls that are the most closely related homeotherm lineage in the multiple alignment data (red). In addition, we defined background homeothermic species (shown in black) in each clade from which we identified conserved regions (CRs, phastCons, FDR < 5%). These background CRs were tested for significant accelerated evolution (ARs, phyloP, Likelihood Ratio Test; FDR < 5%), or deletions in the hibernators and control homeotherms. **(B)** Shows images of each hibernator (blue) and control homeothermic species (red). The numbers of significant accelerated regions (ARs, FDR 5%) and deleted regions (DELs) found in each lineage from the ∼1.3 million background homeotherm CRs tested are shown below the image. The images and data shown correspond to the adjacent species in the phylogenetic tree (A). (**C and D**) The barplots show the numbers of ARs (C) or deleted regions (D) shared in at least 2 of 4 hibernators (parallel hibernator accelerated regions (pHibARs), blue) or homeotherms (parallel homeotherm accelerated regions (pHomeoARs), red). (**E and F**) H3K27ac+ PLAC-Seq reveals significant promoter regulatory contacts genome-wide in the mouse hypothalamus (FDR 1%, min read count >=12, observed/expected > 2; n=10). The heatmap of numbers of contacts shows an example of significant contacts on chromosome 18 (E). The contact plot for *Spats2l* (a significant FFR response DEG) shows how the PLAC-Seq data link pHibARs (light blue track) and pHibDELs (dark blue track) to contacted gene promoters (F). Contact loop height is proportional to the observed/expected read count. Also shown are significant H3K27ac+ ChIP-Seq peaks (FDR 5%, orange tracks), and FFR cycle ATAC-Seq peaks (FDR 5%, black tracks). Gene models are below and promoters are highlighted (grey bars). **(G)** Pie charts of hypothalamus PLAC-Seq contact data show the proportion of pHibARs (light blue) and pHibDELs (dark blue) overlapping 10 kb windows involved in significant promoter-promoter contacts (P-P, darker shade), enhancer-promoter contacts (E-P, medium shade), and those with no significant promoter contacts (i.e. enhancer-enhancer contacts) (No P, light shade). **(H)** The barplot shows the odds ratio of pHibARs (blue) and pHomeoARs (red) occurring in 10 kb regions contacting hypothalamus FFR response DEGs compared to background conserved regions (CRs). pHibARs are significantly enriched for DEG contacting windows compared to pHomeoARs (Woolf test). ****p<0.0001, **p<0.01, *p<0.05. **(I)** We tested pHibARs and pHomeoARs against CRs for enrichment in open chromatin sites (consensus ATAC-Seq peaks, FDR 5% in 4 of 8 replicates, n=8). pHibARs are significantly enriched relative to pHomeoARs (Woolf test). ****p<0.0001; ns, not significant.

To identify genetic elements involved in the evolution of hibernation, we explored convergent genomic changes in hibernators. We tested for parallel hibernator ARs (pHibARs) that impact the same CR in two or more hibernating lineages. We also tested for parallel deleted regions (pHibDELs). We enforced a minimum size of 20 bp for background CR sizes for this analysis but found that peak numbers of pHibARs were observed in ∼40bp CRs (**Fig. S3A**) and in ∼30bp CRs for pHibDELs (**Fig. S3B**). Examples of a pHibAR and pHibDEL are shown in **Fig. S3C and D**. A bootstrap test in which background CRs were randomly sampled without replacement and tested for overlap revealed that significantly more pHibARs and pHibDELs exist in hibernators than expected by chance (**Fig. S3E and F**). Our analysis uncovered 9,374 pHibARs and 25,108 pHibDELs in the mammalian genome (**Fig. 2C,D and Table S4 and S5**). These findings point to convergent evolutionary effects on conserved *cis-*elements.

Distal CREs impact gene expression through long-range 3D interactions. We thus sought to identify genes contacted by pHibARs or pHibDELs in the hypothalamus using H3K27ac targeted proximity-assisted ligation ChIP (PLAC)-Seq (*45*, *46*) (**Fig. 2E,F**). The resulting map of significant contacts allowed us to assign pHibARs, pHibDels and ATAC-seq peaks to contacted gene promoters (**Fig. 2E,F and Table S6 and S7**). We found that 14% of pHibARs reside in distal CREs contacting gene promoters (Enhancer-Promoter (E-P) contacts), 58% make enhancer-enhancer contacts (E-E contacts), and 28% are in CRs without significant regulatory contacts in the hypothalamus (**Fig. 2G**). Similar results were obtained for pHibDELs (**Fig. 2G**).

With this integrated dataset, we probed the interactions between pHibARs and pHibDELs and FFR gene expression responses. We found that pHibARs significantly and disproportionately contact FFR response DEG promoters relative to background CRs. The enriched DEG sets included the 24hr-to-72hr FD and RF period DEG sets (**Fig. S4A**). pHibARs are also significantly over-represented in FFR DEG regulatory contacts compared to control pHomeoARs (**Fig. 2H**). Next, we tested for convergence on active CREs identified from ATAC-Seq peaks and found that pHibARs are significantly enriched in these open chromatin sites compared to both CRs (**Fig. S4B**) and pHomeoARs (**Fig. 2I**). For pHibDELs, we did not find significant enrichments in regions contacting FFR DEGs (**Fig. S4C, E**), but found that pHibDELs are significantly over-represented in ATAC-Seq peaks pointing to active FFR response CREs (**Fig. S4D,F**). Thus, convergent genome evolution in hibernators disproportionately impacts *cis-*regulatory mechanisms involved in hypothalamic FFR responses, pinpointing important CREs and genes.

## pHibAR *Cis-*Elements Target Central Hub Genes Controlling FFR Response Gene Modules

Hub genes are central regulatory nodes governing the behavior and expression of other genes, shaping entire gene co-expression network architectures and molecular programs (*47*). We do not know the nature of the gene co-expression networks for FFR responses in the hypothalamus, the central hub genes involved, or the potential regulatory divergence of these genes in hibernators. Therefore, we first applied weighted gene correlation network analysis (WGCNA) to our RNA-Seq data for the 7 steps of the FFR response to identify gene co-expression modules (**see Fig. 1D**), uncovering 41 modules (**Fig. S5A**). Different modules show different patterns of gene activity across steps of the FFR response (**Fig. S5B**). Gene set enrichment analysis found significant pathway and biological process enrichments for the genes in each module and distinct enrichments for each module that point to different functional roles (**Fig. S6 and S7**). These foundations helped delineate FFR response genetic programs.

To uncover the hub genes for each gene co-expression module, we identified the top 10% of genes ranked according to connection weights to all other genes in a module and then delineated the subset of these that show module membership >0.8 (i.e. correlation to the module eigengene) as hub genes (*48*) (**Fig. 3A and Table S8**). Reactome pathway analysis of the FFR response hub genes identified 73 significant pathways, illuminating hub genes in core metabolic pathways such as organelle biogenesis and maintenance, mitochondrial biogenesis, cellular response to starvation, regulation of apoptosis, MTOR signaling, and other pathways (**Table S8**). We next investigated potential evolutionary and regulatory changes in hibernators.

**Fig. 3.**
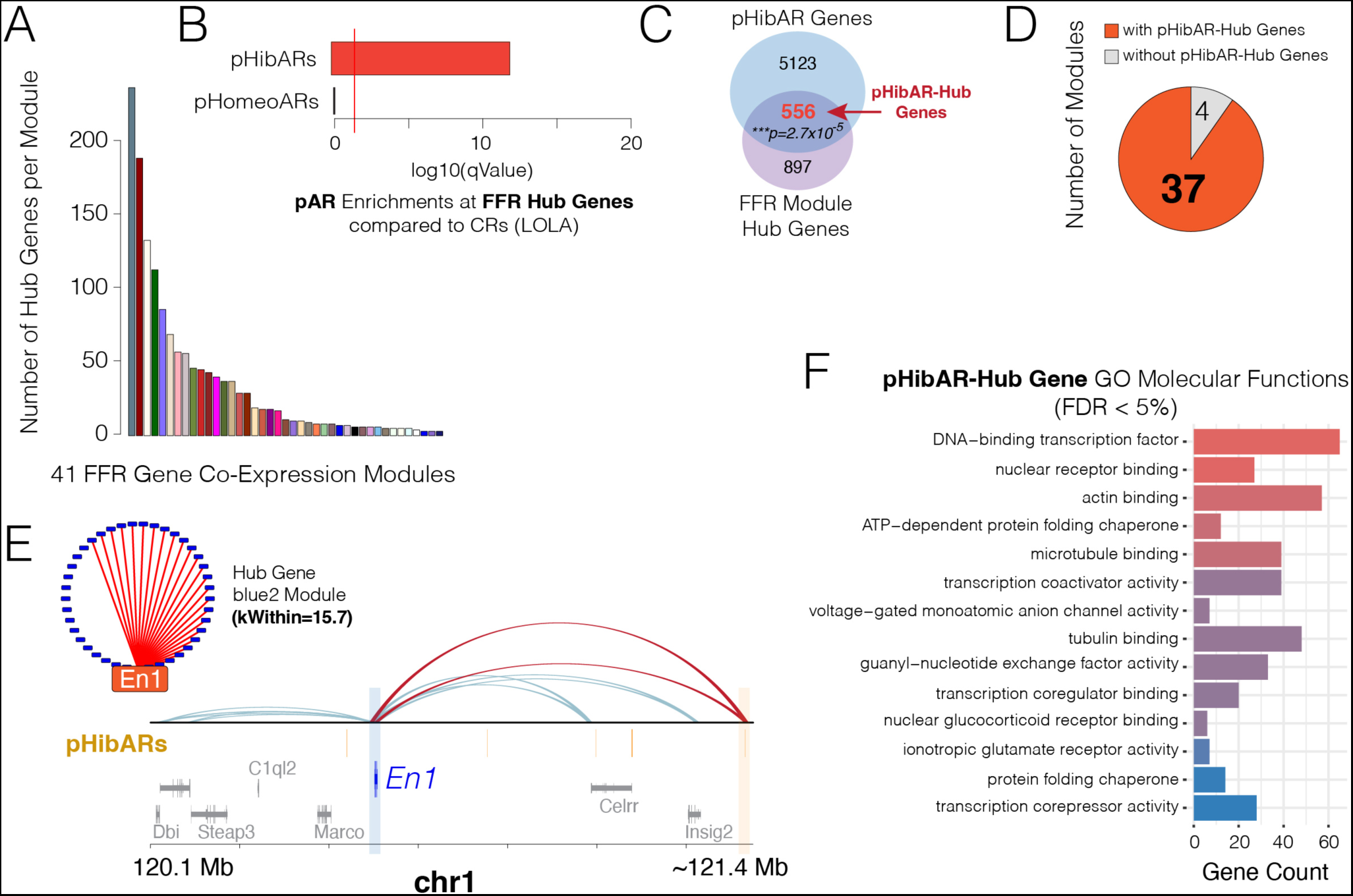
pHibARs regulate hub genes for FFR response gene co-expression modules. **(A)** The barplot shows the number of hub genes identified for each of the 41 FFR response gene co-expression modules uncovered in the mouse hypothalamus. **(B)** The barplot shows the -log10 of the p-value for the enrichment of pHibARs and pHomeoARs at FFR response hub genes compared to background CRs (LOLA Genomic Locus Overlap Enrichment Analysis, R). Red line shows p=0.05. **(C)** The venn diagram analysis shows the number of FFR response hub genes that overlap with genes regulated by CREs with pHibARs (pHibAR-Hub genes), uncovering a statistically significant overlap (Hypergeometric test, p=2.7x10^-5^). **(D)** The pie chart shows the proportion of FFR response co-expression modules that have one or more hub genes regulated by CREs with pHibARs. The results show that nearly all FFR modules have hub genes with sites impacted by convergent genomic changes in hibernators. **(E)** *En1* is a Hub gene uncovered in the blue2 co-expression module linked to dopaminergic neurogenesis (C, see Fig. S5). PLAC-Seq reveals that *En1* is controlled by two regulatory contacts with pHibARs (D, red contacts). The track of pHibARs is shown in orange. Gene models are below. **(F)** The barplot shows the gene ontology molecular function terms that are significant enriched for pHibAR-Hub genes compared to all genes expressed in the adult mouse hypothalamus (FDR < 5%, ClusterCompare R package).

An analysis testing whether pHibARs disproportionately impact some gene modules over others identified 5 modules that are significantly enriched for genes with regulatory contacts from pHibARs (**Fig. S5C**). Gene set analysis showed that the pHibAR enriched modules have roles in neuron differentiation and morphogenesis (**Fig. S5B, darkred module)**; neurotransmitter and opioid signaling (**Fig. S5B, darkolivegreen4 module**); lysosome biogenesis and fatty acid metabolism, and circadian rhythm (**Fig. S5B**, **moccasin module**); inflammatory response and signaling pathways, including Tumor necrosis factor, Receptor Tyrosine Kinase, and Mapk signaling (**Fig. S5B, grey module**); and dopaminergic system function and development (**Fig. S5B, blue module**). Thus, convergent genomic changes in hibernators disproportionately affected a subset of FFR response modules.

By exploring hibernator evolutionary changes to hub genes, we aimed to uncover central regulatory genes impacted by differences between hibernating and homeothermic lineages. Towards this, we found that FFR response hub genes are significantly enriched for regulatory contacts with pHibARs relative to CRs in the mammalian genome (**Fig. 3B**). In contrast, control pHomeoARs do not show a significant enrichment for contacts with FFR hub genes, indicating a hibernator specific effect (**Fig. 3B**). A Venn Diagram analysis found that of all the genes that have regulatory contacts with distal pHibARs, a significant proportion are FFR response hub genes (**Fig. 3C**). We found that 38% of FFR response hub genes have contacts from pHibARs and called this subset ‘pHibAR-Hub’ genes (**Table S8**). Nearly all FFR response modules (37 out of 41) have one or more pHibAR-Hub genes, suggesting hibernation-linked influences on these central regulators exist in most modules (**Fig. 3D**). An example is the homeobox transcription factor, *Engrailed1 (En1),* a top hub gene in the blue2 module that is significantly enriched for dopamine neuron development and function pathways (**Fig. 3E and Fig. S5B, blue module**). The *En1* promoter makes two different regulatory contacts with a distal pHibAR according to our PLAC-Seq data (**Fig. 3E**). In summary, convergent genomic changes in hibernators significantly and uniquely affect *cis-*elements forming regulatory contacts with FFR response hub genes.

A molecular function gene ontology enrichment analysis showed that pHibAR-Hub genes are significantly enriched for regulatory functions, including DNA-binding transcription factor activity, nuclear receptor binding, transcription coactivator activity, corepressor activity, ionotropic glutamate receptor signaling, actin/microtubule/tubulin regulation, nuclear glucocorticoid receptor binding, and more (**Fig. 3F**). Notably, the actin cytoskeleton, microtubules, and intermediate filaments regulate the location, duration, and intensity of signaling in cells (*49*), and glucocorticoid signaling, and the hypothalamic-pituitary-adrenal (HPA) axis that controls adrenal glucocorticoid release, is central to fasting physiological responses (*50*). These findings support the conclusion that pHibAR-Hub genes have regulatory functions that govern the expression of other downstream genes. Thus, we uncovered an initial blueprint of *cis-*regulatory changes and gene expression programs for metabolic control and the divergence of hibernating and homeothermic lineages.

## ARs are Associated with CRE Loss of Function in an Obligate Hibernator

What effects do ARs have on conserved CREs? ARs could arise due to a relaxation of purifying selection. Alternatively, positive selection could drive nucleotide changes in conserved CREs to gain new functionality. We previously predicted that ARs are generally driven by relaxed purifying selection (*25*, *26*). Here, we experimentally tested whether ARs in an obligate hibernator, the 13-lined ground squirrel (squirrel), are associated with differences in hypothalamic gene regulation compared to a homeotherm, namely *Mus musculus,* and associated with loss of function. We performed ChIP-Seq for H3K27ac in the squirrel and mouse hypothalamus to identify active CREs in each lineage and determine whether squirrel ARs are disproportionately in CREs (i) active in mouse and absent in squirrel (mouse-CREs; loss of function), (ii) active in squirrel and absent in mouse (squirrel-CREs; gain of function), or (iii) in CREs active in both species (mouse-squirrel shared CREs) (**Fig. 4A**). Wild squirrels were trapped in summer. Comparative ChIP-Seq profiling uncovered species differences in hypothalamic CRE activity in orthologous genomic regions (**Fig. 4B**) and we identified significant mouse-specific, squirrel-specific, and mouse-squirrel shared H3K27ac+ CRE sites (**Fig. 4C**). The FDR was controlled in each species to define a relatively similar number of significant H3K27ac+ peaks for comparison. Twenty-seven percent of significant H3K27ac+ peaks in squirrel and mouse overlap, indicating shared hypothalamic CREs, while 73% are in nonoverlapping genomic locations indicating species-specific CREs. We tested squirrel ARs for enrichment in species-specific or shared peaks compared to background CRs (**Fig. 4D**). In support of the loss-of-function hypothesis, squirrel ARs are depleted in mouse-squirrel shared CREs and squirrel-specific CREs but significantly enriched in mouse-specific H3K27ac+ CREs (**Fig. 4D,E**). Deletions in background CRs are unambiguous loss-of-function effects on the deleted sequence and we found that squirrel deleted regions are also depleted in shared and squirrel-specific H3K27ac+ peaks CREs but enriched in mouse-specific peaks (**Fig. 4D**). Thus, ARs in an obligate hibernator, the 13-lined ground squirrel, are associated with loss of CREs showing activity in mice.

**Fig. 4.**
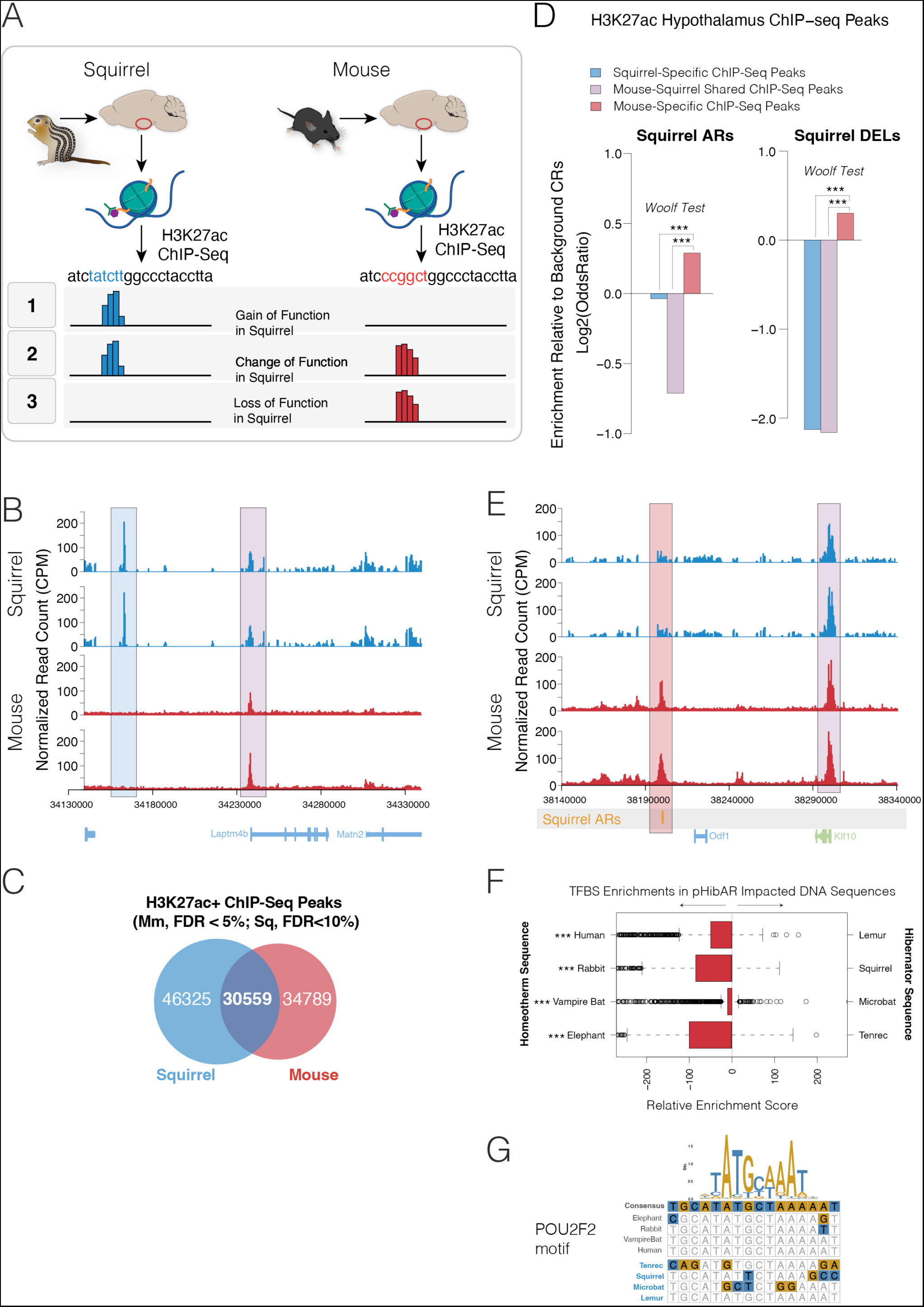
ARs are disproportionately associated with CRE loss-of-function effects in the hypothalamus of an obligate hibernator. **(A)** Schematic summary of potential gain (1), change (2), or loss of function (3) effects of ARs on H3K27ac+ CREs in the hypothalamus of the 13-lined ground squirrel compared to mice. Squirrel ARs in background CRs could be associated with new H3K27ac+ peaks (gain of function in squirrel) compared to mice, loss of H3K27ac+ peaks that are present in mice (loss of function in squirrel) or preservation of H3K27ac+ sites, but a change to DNA sequence motifs (change of function in squirrel). **(B)** Plots show H3K27ac+ ChIP-Seq results for 13-lined ground squirrel (blue) versus mouse (red) adult hypothalamus for two replicates (normalized read counts shown). The results show an example of a species-difference in significant H3K27ac peaks, uncovering a squirrel-specific peak (blue highlight) and squirrel-mouse shared peak (purple highlight). Gene models shown below. **(C)** Venn diagram analysis for the numbers of significant hypothalamus H3K27ac+ peaks for squirrel (Sq) (FDR < 10%) versus mouse (Mm) (FDR < 5%) (n=3). The FDR threshold was adjusted to yield relatively similar numbers of significant peaks for comparison between the two species. **(D)** Barplots show the odds ratio for squirrel ARs and DELs (deleted regions) to overlap with H3K27ac+ peaks that are squirrel-specific (blue), mouse-squirrel shared (purple), and mouse-specific (red) compared to background CRs. Squirrel ARs and DELs are significantly depleted in squirrel-specific and mouse-squirrel shared peaks, but enriched in mouse-specific peaks, which supports a loss-of-function effect for ARs and DELs in the squirrel. Woolf test. ***p<0.0001. **(E)** Plots show H3K27ac+ ChIP-Seq results for 13-lined ground squirrel (blue) versus mouse (red) adult hypothalamus for two replicates (normalized read counts shown). A track indicating the position of a squirrel AR is shown below (orange bar). The results show that an H3K27ac+ peak overlapping the squirrel AR is lost in squirrel but present in mouse. **(F and G)** Boxplot shows differential enrichment score results for TFBS motif enrichments in the orthologous DNA sequences for pHibARs in clade-paired homeotherm versus hibernator species (**F**, PWMEnrich R package, motifDiffEnrichment test; p-value results shown are for a Chi-Square test of observed versus expected counts of enriched TFBS motifs for each species pairing). (**G**) Shows an example TFBS motif in a pHibAR site. ***p<0.001

Given the loss-of-function effects associated with ARs (and deleted regions), we tested whether pHibARs and pHibDELs affect the binding sites for specific transcription factors and regulatory proteins. We found that mouse orthologous sequences for pHibARs are significantly enriched for transcription factor binding site motifs (TFBS) motifs for 44 regulatory proteins (**Fig. S7A**), while sequences impacted by pHibDELs are significantly enriched for motifs for 10 regulatory proteins (**Fig. S7B**). We uncovered top enrichments for motifs for *Nr1h3* (**Fig. S7A**), the *LXRα* nuclear receptor that regulates fat synthesis, cholesterol, and lipoprotein metabolism (*51*). Additionally, motifs for the *E4f1* transcription factor are significantly impacted by pHibARs, which is a key gene regulating cell development, DNA damage responses, and metabolism that is linked to several human diseases (*52*). We tested known TFBS motifs for enrichment within pHibARs for each hibernator relative to homologous sequence for a related background homeotherm. The majority of motifs are significantly enriched in the homeotherm sequences, consistent with a relative loss of TFBS motifs in hibernators (**Fig. 4F and G**). In summary, ARs (and DELs) generally induce loss-of-function in conserved CREs, and pHibARs (and pHibDELs) DNA sequences are relatively depleted for TFBS motifs for defined gene regulatory proteins, suggesting evolutionary modifications that differentiate hibernating and non-hibernating lineages.

## Hibernation-Linked Gene Regulatory Programs Converge on Specific Hypothalamic Cells

The hypothalamus contains an array of cell types that govern metabolic, homeostatic, and behavioral processes. We tested convergent genomic changes in hibernators for effects in specific hypothalamic cell types, potentially linking affected *cis-*elements and genes to molecularly defined cell populations. Therefore, we performed single-cell multi-omic (RNA-Seq + ATAC-Seq) profiling of gene expression and chromatin accessibility in the hypothalamus of fed, 72hr FD, and 12hr RF adult mice. From the RNA-Seq data for 61,400 cells aggregated from all replicates and FFR conditions, we delineated major neuronal and glial cell types (**Fig. 5A**) and uncovered 42 different hypothalamic cell subtype clusters (**Fig. 5B**). Genes that change their expression in specific cells according to FFR conditions (**Fig. S8A**). We tested whether molecular subtypes of hypothalamic cells change in prevalence based on the metabolic state of the animal by tallying the numbers of cells in each subtype cluster according to the FFR condition in which they were detected. Full versus nested mixed-effects models assessed the significance of the interaction between cell subtype and FFR condition. Replicates were treated as random effects. The interaction effect is significant (p=0.0009, Ξ^2^=128, df = 82, LRT), revealing that the prevalence of some molecular cell subtypes depends on the FFR condition of the animal (**Fig. 5C and S8B**).

**Fig. 5.**
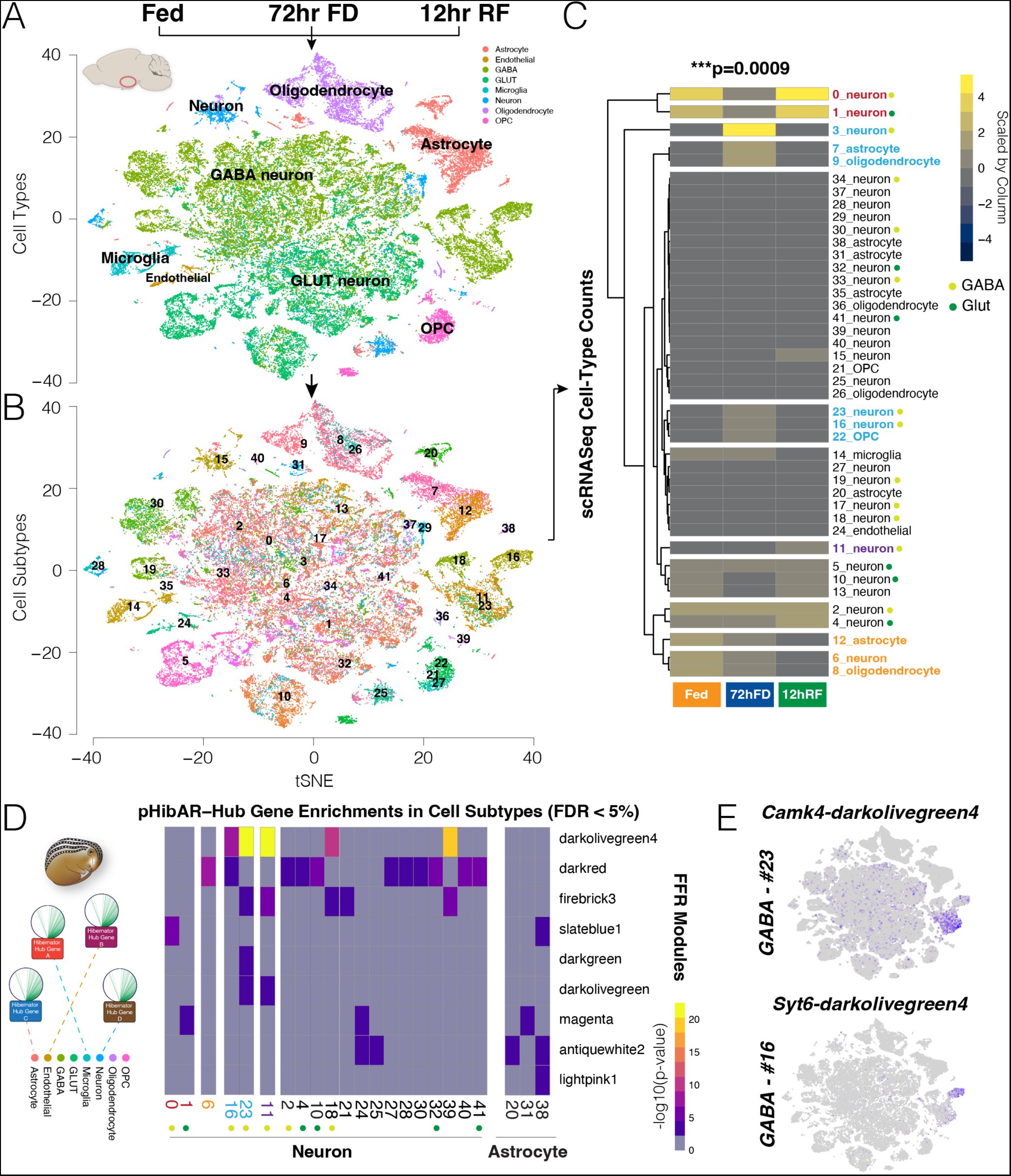
Single-cell multi-omics uncovers FFR alterations to mouse hypothalamic cell types and identifies cell populations disproportionately impacted by pHibAR-Hub genes. **(A and B)** The tSNE plots show the results of Seurat analysis for cell type (A) and subtype (B) clusters from scRNA-Seq hypothalamus data collected for fed (n=2), 72hr FD (n=3), and 12 hr RF (n=3) adult female mice. We identified 8 cell types (A) and 41 cell subtype clusters (B) from 61,400 total cells profiled. **(C)** The heatmap shows the numbers of cells identified from the Fed, 72hr FD, and 12hr RF conditions for each cell subtype (scaled by column). A mixed model testing for an interaction between cell subtype and FFR condition found a significant effect, showing that the numbers of cells detected in each cell subtype depends significantly on the metabolic state (p=0.0009, biological replicates were treated as random effects). Cell subtypes that are enriched (blue) or depleted (red) in the 72 hr FD condition are shown by highlighted text, as well as cell subtypes enriched (purple) or depleted (orange) in the 12hr RF condition. GABA and Glut neuron subtypes are indicated in the legend. **(D)** The heatmap shows the -log10 of p-value for the enrichment of pHibAR-Hub genes from different FFR gene co-expression modules (columns) in different cell subtypes (rows). The results reveal that pHibAR-Hub genes are significantly enriched for expression in subsets of hypothalamic cells compared to all expressed genes in the hypothalamus, including subtypes of neurons and astrocytes. Several cell subtypes (colored labels) that are affected by FFR condition are enriched. GABA and Glut neurons are shown according to the legend in C. **(E)** The t-SNE plots show the expression (purple dots) of the darkolivegreen4 co-expression module pHibAR-Hub genes, *Camk4* and *Syt6,* in hypothalamus cells. The results show *Camk4* is enriched in GABA neuron subtype #23, while *Syt6* is enriched in #16. See panel B.

Six cell subtypes are enriched in the 72hr FD condition (**Fig. 5C, blue text**), such as GABA neuron subtype #3, astrocyte subtype #7, and oligodendrocyte subtype #9; while 2 subtypes are relatively depleted in the 72hr FD versus Fed and 12hr RF conditions (GABA neuron #0 and GLUT neuron #1) (**Fig. 5C, red text**). Additionally, astrocyte, oligodendrocyte, and neuron cell subtypes that are enriched in the Fed state (**Fig. 5C, orange text**), and one GABA neuron subtype enriched in the 12hr RF state (**Fig. 5C, purple text**). Finally, we defined hypothalamic cell subtypes that are stable across FFR conditions, indicating that their molecular identity is not substantially affected by these metabolic changes (**Fig. 5C, black text**). Marker genes delineating each cell subtype are in **Table S9**. Our findings uncovered molecular subtypes of hypothalamic cells, cells that are the most strongly affected by FFR metabolic state, and the genes involved.

We tested whether pHibAR-hub genes are significantly enriched among the marker genes for different cell populations. We found that the pHibAR-Hub genes from 9 different gene co-expression modules are significantly enriched in the marker genes for twenty neuronal and three astrocyte subtypes (**Fig. 5D and Table S10**). Darkolivegreen4 module pHibAR-Hub genes are enriched in GABA neuron subtypes #16, #23, #11, #18 and GLUT subtype #32. Three of these GABA neuron subtypes are affected by metabolic state such that #16 and #23 increase in prevalence during 72hr FD, while #11 increases during 12hr RF (**Fig. 5C**). Indeed, the darkolivegreen4 module exhibits decreased expression during 72hr FD and increased expression during 12hr RF step of the FFR response and is enriched for genes involved in G-protein coupled receptor (GPCR) and opioid signaling (**see Fig. S5B**). *Camk4* and *Syt6* are examples of darkolivegreen4 module pHibAR-Hub genes enriched in GABA neuron subtypes #23 and #16, respectively (**Fig. 5E**). Thus, only some hypothalamic cell types are significantly impacted by pHibAR-hub genes, thereby linking evolutionary genomic changes in hibernators to central hub genes for FFR responses to discrete hypothalamic cell-types.

## Chromatin Remodeling Response to FFR Changes Varies Across Hypothalamic Cells

We next focused on the activity of individual *cis-*elements at the cellular level. Having found that pHibARs are enriched in accessible chromatin sites identified from bulk ATAC-Seq, we elucidated cellular chromatin accessibility and dynamics. Building on our bulk tissue analysis showing that RF is a critical period of chromatin changes, single cell multi-omics profiling of FFR conditions revealed that RF induced significantly increased numbers of accessible chromatin sites in astrocytes, endothelial cells, GABA neurons, and oligodendrocytes, but not in GLUT neurons, microglia, or other neurons (**Fig. 6A,B**). The 72-hr FD condition increased chromatin accessibility sites in astrocytes, endothelial cells, and GABA neurons compared to the Fed condition (**Fig. 6A**). Many distal ATAC-Seq peaks pointed to putative CREs not yet annotated by ENCODE (*53*), especially those uncovered in the RF condition (**Fig. S11**). These results demonstrate important metabolic effects on CREs at the cellular level, reveal how FFR states differentially affect chromatin according to hypothalamic cell type, and uncover putative active CREs for each cell type and FFR metabolic state.

**Fig. 6.**
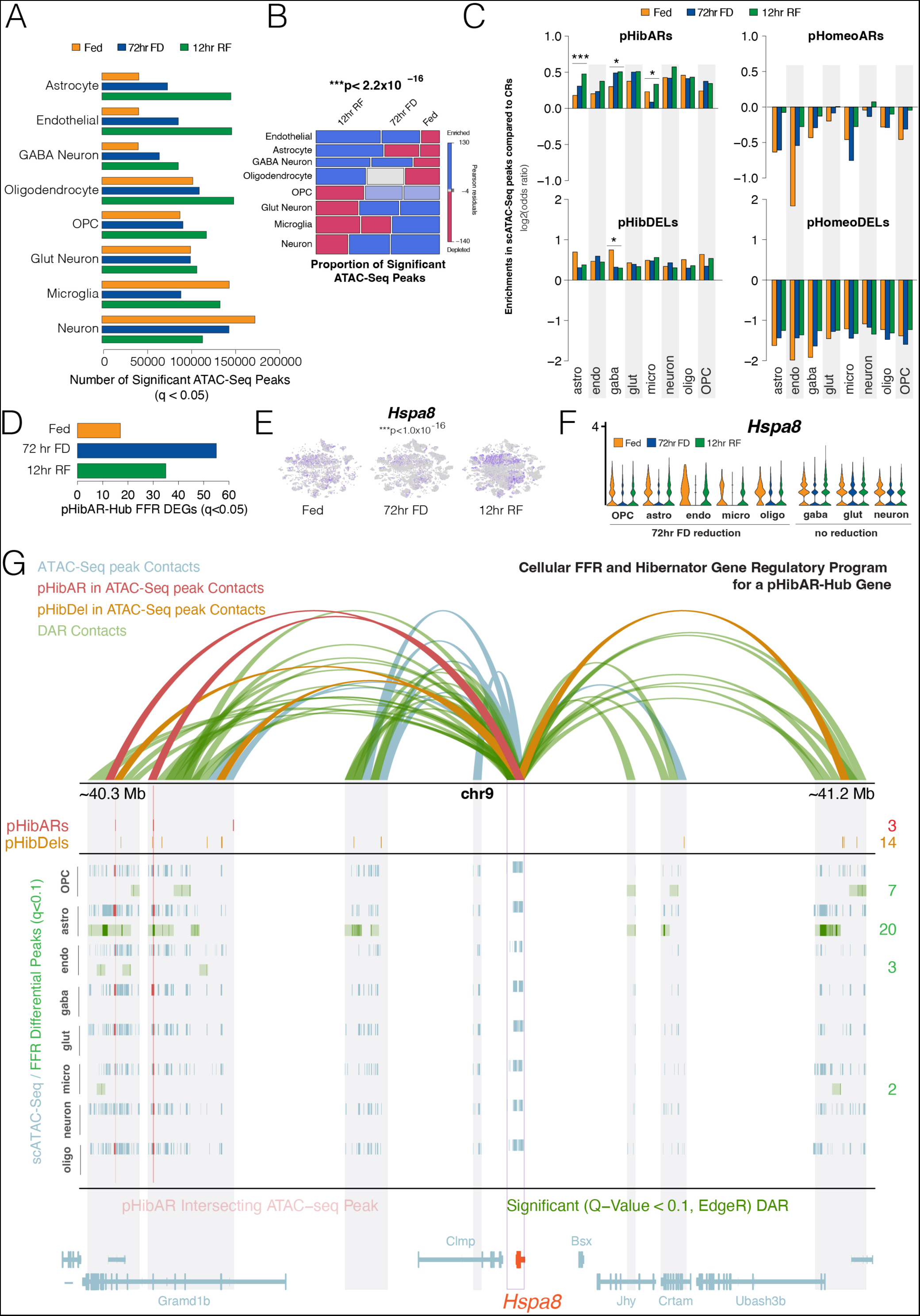
FFR responses cause cell-type dependent changes to chromatin accessibility in the hypothalamus and reveal hibernation-linked genetic programs at the cellular level. **(A)** The barplot shows the number of significant ATAC-Seq peaks (FDR < 5%) uncovered for each hypothalamic cell type and FFR condition. The results reveal 72hr FD and 12hr RF associated increases in the numbers of accessible chromatin sites in astrocytes, endothelial cells, GABA neurons, oligodendrocytes, and OPCs, but not other cell types. **(B)** Mosaic plot analysis of numbers of ATAC-Seq peaks according to cell type and FFR condition reveals a significant dependency (Chi-Square p-value shown), showing that chromatin accessibility is significantly enriched during 12hr RF in endothelial cells, astrocytes, GABA neurons, and oligodendrocytes. Pearson residuals show relative enrichment (blue) versus depletion (red) effects for each category. **(C)** The barplots show the odds ratio for pHibARs, pHomeoARs, pHibDELs, and pHomeoDELs to be located in open chromatin sites in hypothalamic cell types in Fed, 72hr FD, and 12hr RF conditions compared to background CRs. pHibARs and pHibDELs are enriched (p<0.05 for all conditions), while pHomeoARs and pHomeoDELs are depleted. A comparison between FFR conditions for pHibARs showed FFR dependent differences in enrichments for astrocytes, GABA neurons, and microglia, and in GABA neurons for pHibDELs (Woolf test). ***p<0.001, *p<0.05 **(D)** Barplot showing the numbers pHibAR-Hub genes showing significant differential expression at the cellular level according to FFR condition (FDR < 5%). **(E)** scRNA-Seq tSNE plots display *Hspa8* expression across FFR conditions, showing decreased expression in the 72hr FD condition. A model comparing *Hspa8* expression summed from all cells in the 72hr FD condition to the Fed and 12hr RF conditions found a significant difference (p<1.0x10^-16^, Seurat). **(F)** Violin plots show pHibAR-Hub gene, *Hspa8,* expression across cell types and FFR conditions. Expression is reduced in the 72hr FD condition for OPCs, astrocytes, endothelial cells, microglia, oligodendrocytes, and GABA neurons, but not strongly affected in glutamatergic (Glut) and other neurons (neurons). **(G)** A hibernation-linked gene regulatory program for *Hspa8* in hypothalamic cells. The H3K27ac+ PLAC-Seq data shows *Hspa8* promoter contacts in the adult mouse hypothalamus (10kb cis-bin windows). Tracks below show pHibARs (red), pHibDELs (orange), and significant ATAC-Seq open chromatin peaks (blue) for each cell type. ATAC-Seq peaks that overlap with pHibARs are red and highlighted by red lines. The green tracks show ATAC-Seq peaks detected with significant differential chromatin accessibility between Fed, 72hr FD, and 12hr RF conditions for each cell type (q<0.1) and the number of detected sites is shown on the right (green text). PLAC-Seq bins overlapping differentially accessible chromatin regions (DARs) are highlighted in green. FFR effects on chromatin accessibility for *Hspa8* CREs are strongest in astrocytes, OPCs, endothelial cells, and microglia. *Hspa8* regulatory promoter contacts are also colored according to whether the distal contact overlaps DARs and/or convergent genomic changes in hibernators (pHibARs or pHibDELs) as follows: FFR response DARs (green), pHibARs in ATAC-Seq peaks (red), pHibDELs in ATAC-Seq peaks (orange), and contacts overlapping other significant ATAC-Seq peaks (blue). Gene models are shown below. Purple box shows promoter contact sites associated with *Hspa8* (orange gene model).

We determined how convergent genomic changes in hibernators impacted accessible *cis-*elements for different cell types and FFR states. Our analysis revealed that both pHibARs and pHibDELs are significantly over-represented in cellular ATAC-Seq peak sites for all 8 cell types compared to background CRs (**Fig. 6C**). In contrast, pHomeoARs and pHomeoDELs show the opposite and are under-represented in these sites (**Fig. 6D**). Thus, genomic changes in hibernators impact active CREs in all major hypothalamic cell types, yet the magnitude of the enrichment effect is significantly influenced by the FFR state and cell-type. For astrocytes, GABA neurons, and microglia, we found significantly greater pHibAR enrichments for distal ATAC-Seq peaks detected in the 72 hr FD and RF conditions compared to the fed condition (**Fig. 6C**). pHibDELs are significantly more enriched in CREs detected in GABA neurons in the fed condition versus the 72hr FD and RF conditions (**Fig. 6C**). Thus, *cis-*element evolutionary changes in hibernators disproportionately intersected with CREs active in specific hypothalamic cell types in specific metabolic states. Our cellular analysis pinpoints putative CREs important in cellular FFR responses and evolutionary divergence (**Data S1-3**).

## Cellular Genetic Programs for Hypothalamic FFR Responses and Hibernator Evolution

In a final analysis, we aimed to elucidate a coherent framework of cellular hypothalamic *cis-*elements and programs underlying FFR responses, and their modification in hibernating species. To begin, we identified the pHibAR-Hub genes with significant gene expression changes at the cellular level according to FFR state. This uncovered 17, 55, and 35 pHibAR-Hub genes with significant differential expression in the fed, 72hr FD, and 12hr RF conditions, respectively, relative to each other (**Fig. 6D**). Some pHibAR-Hub genes increase or decrease their expression in a particular cell type according to metabolic state. *Hspa8* is a member of the heat-shock protein 70 (HSP 70) family with roles in autophagy and protein-folding that breaks down functional amyloids in the brain (*54*), and has important roles in neuroprotective processes and effects on protein misfolding in neurodegenerative diseases (*55–61*). We found that *Hspa8* expression is reduced in the 72hr FD condition in glia and GABA neurons, but not in glutamatergic or other neurons (**Fig. 6E, F**). Changes to *Hspa8* expression can have neuroprotective effects (*54*, *60*) and influence the accumulation of protein aggregates involved in neurodegenerative processes (*62*). Importantly, hibernators evolved mechanisms for enhanced neuroprotection and resolving tauopathy (*14*), and we therefore performed a detailed analysis to understand the regulatory program of this candidate gene at the cellular level.

Integration of our PLAC-Seq, single cell ATAC-Seq, and hibernator convergent genomics results revealed a cellular regulatory program governing *Hspa8* expression (**Fig. 6G**). The *Hspa8* promoter contacts with CREs showing open chromatin in different cell types. We found CREs that show significant differential accessibility between FFR states and determined that CREs in astrocytes, OPCs, endothelial cells, and microglia are significantly affected, but not those in oligodendrocytes, GABA neurons, Glut neurons, or other neurons (**Fig. 6G**). These findings generally agree with our scRNA-Seq data showing cell type dependent FFR changes in *Hspa8* expression (**Fig. 6E**). FFR effects on chromatin accessibility impacted the largest number of CREs in astrocytes and revealed an active CRE in multiple cell types (significant ATAC-Seq peak), a pHibAR, and shows FFR differential accessibility in astrocytes (**Fig. 6G, purple contact**). Thus, the intersection of different evolutionary and genomics datasets unveils cellular *cis-*regulatory programs for FFR responses, which we uncovered for each of the pHibAR-Hub genes in the hypothalamus (**Data S1-3**).

## Discussion

We have elucidated *cis-*regulatory programs within the hypothalamus that underpin mammalian metabolic and molecular responses to food scarcity and the genomic divergence of hibernating and homeothermic lineages. Our results build on the pivotal role of the hypothalamus in hibernation and evolutionary changes to energy homeostasis to reveal CREs, genes, pathways, and cell types that orchestrate these physiological processes. Multi-omics datasets profiling FFR genomic responses in mice were integrated with a comparative genomics analysis of hibernating versus homeothermic mammals that identified shared *cis-*regulatory changes in species that independently evolved obligate hibernation. Convergent evolution in hibernators was revealed to target conserved *cis-*elements governing hypothalamic FFR responses. Hub genes that are central regulatory nodes of FFR responses are disproportionately affected by convergent genomic changes in hibernators, but not homeotherms, uncovering a core framework of metabolic and hibernation control that helps to prioritize genetic targets and hypothalamic cell types for study. Our genetic blueprint for mammalian metabolic control offers a unique vantage point for now testing how metabolic setpoints, obesity-related diseases, neurodegenerative processes, cell protection, and aging can be genetically and epigenetically reprogrammed, a concept that holds promise for tackling disease and is supported by functional studies in our companion paper (*38*).

Hibernation is an adaptation to extended periods of food scarcity with potential applications in medicine and space travel (*63*, *64*). The hypothalamus is a central controller of metabolism and physiology, harboring neural circuits that can induce torpor-like states in mice (*65–67*). We found that convergent genomic changes in obligate hibernators, including parallel ARs and deletions, are overrepresented in conserved *cis-*elements regulating hypothalamic gene expression. By studying mice, which are capable of brief fasting-induced torpor bouts, we determined that these changes disproportionately accumulated at CREs for genes that change expression during the transition from a fasting to a starvation response, and during the refeeding period after food deprivation-induced torpor. We found that chromatin remodeling and gene expression changes are especially prevalent during refeeding, affecting astrocytes, endothelial cells, oligodendrocytes, and GABAergic neurons. Convergent genomic changes in hibernators point to CREs with roles these refeeding responses and divergence in hibernators. Although the refeeding period is not as widely studied as torpor or fasting, it is a critical period of healing and recovery with biomedical significance. Following extended torpor, hibernators must rapidly recover from immobility (*15*, *68*, *69*), nutrient deprivation, insulin resistance (*16*, *70*), accumulated wastes and toxins (*71*), and tauopathy (*14*). Moreover, synaptic connections are regenerated, and neuroprotective mechanisms defend against brain damage from reperfusion with suddenly increased blood flow that has similarities to stroke (*11*). Our study illuminates CREs and cellular mechanisms involved.

Hibernators evolved unique capabilities for longevity, neuroprotection, synapse regeneration, cell damage prevention, and tauopathy resolution (*13*, *14*, *19*, *72*, *73*). Intermittent fasting, metabolic changes, and torpor in mice can rescue neurodegeneration (*12*, *39*, *74*). We found that modules of co-expressed genes across different FFR states in the mouse hypothalamus unveiled hub genes governing FFR responses, and convergent genomic changes in hibernators significantly accumulated at CREs controlling these hub genes. These pHibAR-Hub genes are overrepresented in pathways involved in neurodegeneration process pathways. Observed enrichments in mitochondrial functions, protein-folding, protein ubiquitination and catabolism, and stress-responses are not only central to fasting biology, but also key mechanisms in neurodegenerative diseases (*1*). Thus, pHibAR-Hub genes include important candidate target genes for affecting processes involved in neurodegeneration and brain aging.

Hibernators become seasonally obese, showing metabolic and foraging adaptations that support binging on high-calorie foods for fat accumulation before hibernation, then becoming insulin resistant and switching to fat metabolism during prolonged winter torpor (*7*, *75*). ARs and deletions in a model hibernator – the 13-lined ground squirrel – are predominantly associated with loss-of-function effects at CREs conserved in homeotherms and active in the hypothalamus. This suggests that pHibARs and pHibDELs point to CREs that are integral to homeothermic metabolic and behavioral regulation, but unnecessary for hibernators or impede capabilities for fat accumulation, heterothermy, torpor, metabolic suppression, or other critical hibernator traits. Our single cell multi-omics results show the cellular activity of these *cis-*elements. In our companion study, deletions of individual CREs with pHibARs in mice altered gene expression, metabolism, weight gain on an obesogenic diet, and foraging (*38*). This supports function roles for CREs illuminated by pHibARs and pHibDELs, providing a genetic framework for the regulation of metabolism and evolution of hibernation.

## Acknowledgments

Thank you to Drs. Amandine Chaix, Jared Rutter, Scott Summers, and all members of the Gregg lab for commenting on the manuscript. The CLAMS metabolic phenotyping experiments were performed with the University of Utah Metabolic Phenotyping Core Facility. Bulk and single cell genomics experiments and sequencing were performed with the University of Utah High Throughput Genomics Core. Squirrel tissue was obtained from Dr. Dana Merriman (Squirrel Colony at the University of Wisconsin Oshkosh). Thank you to Tyler C. Leydsman and Bennett Cottle for technical and mouse colony support.

## Author contributions

Conceptualization: EF, ACR, JGM, CG

Methodology: EF, JGM, ACR, SS

Investigation: EF, JGM, ACR, SS, DT, CSH, PJMC, CG

Visualization: EF, CSH, JGM, CG

Funding acquisition: CG

Project administration: CG

Supervision: CG

Writing – original draft: CG

Writing – review & editing: EF, JGM, ACR, SS, DT, CSH, PJMC, CG

## Data and materials availability

All data are available in the main text or the supplementary materials, and all genomics files are deposited in the NIH Short Read Archive repository.

## Supplementary Materials

Materials and Methods

Figs. S1 to S11

Tables S1 to S10

Data S1 to S3

