## Supplementary Materials for "Genomic Convergence in Hibernating Mammals Elucidates the Genetics of Metabolic Regulation in the Hypothalamus"

**The PDF file includes:**

Materials and Methods
Figs. S1 to S11
Tables S1 to S10
Data S1 to S3
References

### Materials and Methods

### EXPERIMENTAL MODEL AND SUBJECT DETAILS

*Mice*

**Housing and Husbandry**

All experiments were conducted in compliance with protocols approved by the University of Utah institutional animal care and use committee (IACUC). Mice were bred and housed on ventilated racks at the University of Utah Comparative Medicine Center on a 12hr light cycle, 6am on and 6pm off, with an ambient temperature range of 20-25C. All mice were given water and food (Harlan-Teklad 2920X soy protein-free) ad libitum, with the exception of fasting periods for genomic and torpor experiments. Adult breeders (6-weeks to 1-year of age) were paired continuously, and pups were weaned at postnatal day 21 (P21) and cohoused with up to five same-sex littermates or similar aged same-sex mice of the same line; mice were never singly housed. Before dissections of brain tissue, mice were put to sleep with isoflurane gas anesthesia and decapitated.

**C57Bl/6J Mice**

C57Bl/6J mice for genomics experiments were commercially acquired from Jackson Labs (000664). CASTEiJ mice for generating C57Bl/6J x CASTEiJ F1 hybrid mice for H3K27ac mus musculus PLAC-Seq were acquired from Jackson Labs (000928). F1 hybrid mice were generated from reciprocal C57Bl/6J x CASTEiJ matings.

**Wild-Caught 13-Lined Ground Squirrel**

Freshly-dissected adult female flash-frozen hypothalami were obtained from the Squirrel Colony at the University of Wisconsin Oshkosh under Dr. Dana Merriman's direction. Squirrels were trapped and harvested for tissue in July and August.

### METHOD DETAILS

<u>Bulk Tissue Genomics</u>

**RNA-seq**

Hypothalamus was dissected from adult female mice midday and RNA was isolated using the RNeasy Lipid Tissue kit (Qiagen 74804), treated with DNAse (Qiagen 79254). For food deprived mice, mice were moved into a clean cage and food was removed for 24-hrs, 72-hrs, or 72-hrs with reduced ambient temperature (18°C) prior to harvesting, with similar circadian timing to fed mice (midday). Refed mice were exposed to 72-hrs FD and then provided ad libitum access to mouse chow for 1hr (1hr RF), 12hrs (12hr RF), or 1 week (1 week RF). RNA-seq was performed by the Huntsman Cancer Institute High-Throughput Genomics Core at the University of Utah, using NEBNext Ultra II Directional RNA Library Prep with rRNA Depletion Kit (human, mouse, rat) library prep and NovaSeq S4 Reagent Kit v1.5.

**ATAC-seq**

At eight weeks of age, we dissected, and flash froze female mouse brain hypothalamic tissue for Fed, 72-hr FD, 12-hr RF, and 1-week RF (n=8). Samples were prepared for ATAC-Seq profiling of open chromatin using a commercial ATAC-Seq Kit (Active Motif, #53150).

**PLAC-seq**
8-10 week old adult female C57Bl6/J x CasteEiJ F1 hybrid mouse hypothalamus was dissected and flash-frozen (n=10). A ChIP-grade antibody for H3K27ac (AbFlex Histone H3K27ac rAb, Active Motif, cat #91193) was used. Samples were prepared for PLAC-Seq profiling of regulatory contacts using a commercial kit (Arima Hi-C kit+, Arima Genomics). Libraries were prepared for sequencing by the University of Utah High Throughput Genomics core facility using the NEBNext ChIP-Seq Library Prep Reagent Set for Illumina (E6200). Sequencing was performed using an Illumina NovaSeq 6000 with NovaSeq S1 Reagent Kit v1.5_150x150 bp paired end sequencing.

**H3K27ac ChIP-Seq**
H3K27ac 8-10 week old adult female mouse or adult female 13-lined ground squirrel hypothalamus was dissected and flash-frozen. A ChIP-grade antibody for H3K27ac (AbFlex Histone H3K27ac rAb, Active Motif, cat #91193) was used for chromatin immunoprecipitation. ChIP was performed using the ChIP-IT High Sensitivity kit (Active Motif, Cat# 53040). Libraries were prepared for sequencing by the University of Utah High Throughput Genomics core facility using the NEBNext ChIP-Seq Library Prep Reagent Set for Illumina (E6200). Sequencing was performed using an Illumina NovaSeq 6000 with NovaSeq S1 Reagent Kit v1.5_150x150 bp paired end sequencing.

<u>Single cell Multi-Omics</u>

**Hypothalamus single nucleus RNA-Seq + ATAC-Seq**
At eight weeks of age, we collected female mouse brain hypothalamic tissue for Fed (n=2), 72hr FD (n=3), and 12hr RF(n=3) conditions. We flash froze the tissue in liquid nitrogen and stored in -80 °C freezer. We follow the 10X GENOMICS nuclei isolation protocol specific for single cell multiome ATAC + Gene Expression (CG000505, PN-1000493, PN-1000494). In brief, tissue was submerged in 200 uL of Lysis Buffer and dissociated with pestle and incubate in ice for 10 minutes. Passing through a column, we centrifuge the dissociated tissue at 16,000 rcf for 20 seconds at 4 °C. Then, we discarded the column, vortexed the flow and centrifuge for additional 3 minutes at 500 rcf and 4 °C. We removed the supernatant, resuspended the pellet in 500 uL Debris Removal Buffer, and centrifuged at 700 rcf for 10 minutes at 4 °C. Finally, we discarded the supernatant, resuspended the nuclei pellet in 30 to 50 uL of Resuspension Buffer and proceed to calculate nuclei concentration, nuclei quality assessment, library preparation, and sequencing. We prepared all buffer following the vendor’s protocol.

The CellRanger - arc version 2.0.2 pipeline (10X Genomics) was used to process the scRNA-seq and scATAC-seq FASTQC files. The cellranger counts command was used to align the raw FASTQ files to the mm10mouse reference genome. For gene expression sequencing, filtered count matrices were analyzed using the R packager Seurat (V5.0.0) [1]. We emphasized in the designed for the integration of single-cell RNA sequencing (scRNA-Seq) datasets, specifically targeting the correction of batch effects in all replicates. Samples were merged into a single

Seurat object to perform consistent filtering and quality-control (QC). Cells with low features or unique molecular identifier (UMI) counts or high mitochondrial read percentage were discriminated. In addition, to remove doublets, cells with abnormally high UMI counts were removed. Finally, any cells expressing mutually exclusive markers were removed. At the end, 61,400 cells passed filtering and QC processing. Data normalization was done using SCTransform R package, and dimensionality reduction techniques, including PCA and t-SNE, were applied. The data integration to correct batch involved the identification of integration features, anchors, and subsequent data integration using Canonical Correlation Analysis (CCA). The dimensional reduction t-SNE is applied uncovering 42 different hypothalamic cell subtype clusters.

Using the conserved markers function in Seurat and Celldex (v1.13) R packages, we identified eight different cell types: Astrocytes, Endothelial, GABA neurons, Glutaminergic neurons, Microglia, Other Neurons, Oligodendrocytes, and Oligodendrocyte Precursor Cells (OPC) and assigned their identity to the corresponded cluster. Consequently, we performed gene expression analysis using the conserved marker function of the Seurat package at between the different metabolic categories (FFR) at the cell level and cluster level.

<u>Metabolic Phenotyping</u>

**Body Composition and Weight**

For torpor experiments, body weight and composition were measured (NMR, Bruker Minispec LF50) prior to logger implantation and after logger removal post-CLAMS experiment. Body weight was also measured before mice were placed into CLAMS.

**Surgical Implantation of Body Temperature Logger**

At 2.5-5 month of age and body mass range of 18-25g, genotypically-balanced cohorts of pHibAR KO and WT female mice were surgically implanted with a temperature data logger (Star-Oddi, DST nano-T) programmed to record at 10-minute intervals. The mice were anesthetized with an isoflurane vaporizer, administered with lidocaine (0.003mg/g) and carprofen (0.005mg/g), and the logger was subcutaneously implanted through a 1cm mid-scapular incision and closed with surgical wound clips. Mice were administered with analgesics for two days post-op and singly housed for at least two weeks to recover, with food and water available *ad libitum*.

**Torpor Induction and Metabolic Monitoring**

The mice with body temp implants were singly housed in metabolic CLAMS cages (Comprehensive Lab Animal Monitoring System, Columbus Instruments; University of Utah Metabolic Phenotyping Core) for total one week. Mice were allowed to acclimate to the cage, with regular food, at 25C for the first two days. On day 3 at 12pm, food was removed and the ambient temperature was dropped to 18C to induce torpor behavior. After 48hr, mice were refed and warmed back to 25C and monitored for another 72hr. At the end of the experiment, mice were euthanized for retrieval of the body temperature loggers and body weight and composition were again measured.

<u>Defining Accelerated Regions</u>

We subset and reformatted the Zoonomia 241-mammal CACTUS alignment (MAF file) using grep commands, MafFilter, and GNU parallel to include only the species of intersect (Fig 2A). We then used the command phastCons from the R package rphast to define regions conserved in the 8 background species (Fig 2A, parameters rho=0.4, coverage=0.2, expected.length=40). (Hubisz et.al. 2011) We required the presence of a minimum of two hibernator sequences and excluded coding regions. We used the rphast function phypoP to test each conserved region (CR) for acceleration in each hibernating species, calculating socres with the likelihood-ratio test. (phyloP parameters: mode = "ACC", method = "LRT"). We used the rphast function msa.sample to randomize the conserved sequences and test for acceleration in the randomized sequences, allowing us to compare acceleration scores and calculate p-values for the original sequences (Hubisz et.al. 2011). False discovery rates were calculated for the 1,255,387 autosomal CRs with the R package qvalue. We designated regions with significant acceleration (q-value < 0.05) in 2 or more of the 4 hibernating species pHibARs (parallel hibernator accelerated regions). CRs were defined in human coordinates (GRCh38) and lifted-over into mouse coordinates (GRCm38) for comparisons with other mouse data. Regions longer than 1000 bp after liftover were discarded. We followed the same protocol to define homeotherm accelerated regions for 4 homeothermic species against the same 8 background species. We designated regions significantly accelerated (q-value < 0.05) in 2 or of the 4 homeothermic species pHomeoARs (parallel homotherm accelerated regions).

Defining Deleted Regions

We defined regions conserved in the 8 background species using the MAF file generated for the accelerated regions described above with phastsCons (Fig 2A, parameters: rho=0.4, coverage=0.2, expected.length=40). We used the rphast function informative.regions.msa to test the background sequences for presence in the alignment (parameters: gaps.inf = F). We included only CRs present in all 8 background species. We excluded coding regions from our search. We used the rphast function informative.regions.msa to define regions of missing sequence form the multi alignment. CRs containing cumulative deletions longer than 10 bp in 2 or more hibernating species were designated as pHibDels (parallel hibernator deletions.). CRs were defined in human coordinates (GRCh38) and lifted-over into mouse coordinates (GRCm38) for comparisons with other mouse data. We used the same protocol to define pHomeoDels (parallel homeotherm deletions).

### QUANTIFICATION AND STATISTICAL ANALYSIS

Bulk RNA-seq

We used hisat2 python scripts hisat2_extract_splice_sites.py, hisat2_extract_exons.py and the command hisat2-build to build GRCm38 transcriptome index. We aligned paired-end RNA-seq reads to this transcriptome index with the sequence aligner hisat2. We used featureCounts from rsubread to count reads in genes in a gtf file downloaded from ensemble.org for the mouse genome build GRCm38 (release 102). Only protein coding genes were considered in our differential expression analyses performed with EdgeR. False discovery rates were calculated with the R packages 'qvalue'.

Bulk PLAC-seq

PLAC-seq reads were assessed for quality with MAPS, combined and processed with the MAPS pipeline (Juric, et.al. 2019). Significant contacts were designated with the following parameters and thresholds: 10 kb interaction windows, ≤ 1 MB contact range, < 1% FDR, mapq ≥ 30, ≥ 2-fold enrichment, and ≥ 12 read depth minimum (Juric, et.al. 2019). Output BEDPE files were intersected with a promoter regions BED file downloaded with the UCSC table browser indicating regions 2 kb upstream from each transcription start site with bedtools pairtobed (parameters: -type either) to identify potential promoter region contacts. We used custom R scripts to mark PLAC-seq contact bins as promoter intersecting bins and distal bins. The resulting contact tables were used to assign ATAC-seq peaks, ChIP-seq peaks, CRs and ARs to promoter genes.

H3K27ac ChIP-Seq
Mouse and squirrel H3K27ac ChIP-seq peaks were called with the peak caller macs2. It was important for our analysis to compare similar numbers of ChIP-seq peaks for the two species, even at the expense of some peak confidence. We used the intersect of the two mouse H3K27ac peak sets (Q-Value < 0.05) and the union of the two squirrel peak sets (Q-Value < 0.10). We used bedtools intersect (parameters: -wa ) to calculate the overlaps between the two H3K27ac peak sets. We designated mouse H3K27ac peaks that did not intersect squirrel peaks as 'mouse-specific' and mouse peaks that did intersect as 'shared' while unintersecting squirrel peaks were counted and 'squirrel specific'. These peak sets were tested for enrichment with squirrel ARs and deletions relative to CRs with a Fisher's Exact Test. AR and deletions enrichments were tested for independence with the Woolf Test implemented in the R package vcd (4D).

TFBS motif Enrichment Analysis
The human DNA-sequences of ARs, deletion containing regions and CRs were extracted with bedtools getfasta. We tested AR and deleted region sequences against CR sequences for enrichments with human motif sets (HOCOMOCO v11 CORE) with the meme suites SEA tool. For differential motif enrichments in pHibAR orthologous sequence for hibernator-homeotherm paired species, we used the function motifDiffEnrichment() from the the R package PWMEnrich to test enrichment of known motifs in pHibAR sequences from each of the 4 hibernating species against homologous sequence from a related background homeotherm. We tested 2,287 motifs from the R package PWMEnrich.Hsapiens.background using the 'logan' background correction with human sequence. Statistical testing comparing the counts of numbers of enriched TFBS motifs between each species pairing was performed using a Chi-Square test of observed vs expected counts.

Single cell Multi-Omics
scATAC-seq alignment files were divided by cell type with subset-bam (https://github.com/10XGenomics/subset-bam) using barcodes determined by**. . .(Josue)** ATAC-seq peaks were called for the 8 cell types and 3 FFR conditions with the peak caller Genrich using the following parameters: -r -j -l 140 -d 100 -E ENCFF547MET.bed.gz where ENCFF547MET.bed.gz is a BED file of 'exclusion regions' provided by ENCODE: https://www.encodeproject.org/files/ENCFF547MET/ We tested these peaks for enrichment with pHibARs, pHomeoARs, pHibDels and pHomeoDels with the R package LOLA.

To define differentially accessible regions (DARs) for all cell types, for each cell type we took the union of the scATAC-seq peaks across FFR samples. We counted ATAC-seq reads in these union peaks with featureCounts from subread (v1.5.3) with the following parameters: -F SAF -p -B -C -M --largestOverlap --primary -T 24 -Q 30. We parallelized these commands with GNU parallel. For each cell type we used EdgeR to test the counts for a main effect of treatment. We adjusted for multiple testing with the rpackage qvalue. (Storey Paper). q-values < 0.10 were considered significant.

Enrichments of ARs relative to CRs in scATAC-seq peaks were calculated with the R package LOLA.

**Contact plots**
Contacts plots were created with the R package plotgardener. We added 200 bp to either end of each AR to make them more visable.

<u>Metabolic Phenotyping</u>

**CLAMS data graphing**
All CLAMS data from the torpor experiments were exported cropped to start at 6pm of the first day when the first light cycle starts and end at 12pm of the last day. Data was initially analyzed in CalR and hourly average files were exported for further analysis in R. Body temperature collected through implants and CalR data were combined. Each measure was visualized as hourly averaged as line graphs with SEM using the plotly package in R.

**Generalized Linear Modeling of CLAMS data**
For the statistical analysis of the torpor assay, we used the CalR and Body temperature data as 6 hour averages. Due to some very high measurements for Respiratory Exchange Ratio (RER) as levels that are physiologically impossible, we set all values higher than 1.7 to 1.
Food consumption data was set to a maximum of 3.1 kcal/hr as this represents 1g of food consumed by the mouse in 1 hr. This is assumed to be the maximum food intake possible for a mouse.

For each light or dark phase, 2 6-hr periods were combined. Generalized linear modeling was performed using a Gaussian distribution to test main and interaction effects with a likelihood ratio test for full and nested models.

For the 1 week refeeding data, we compared the entire Fed phase to the Refeeding phase or Day 1 (Fed) vs Day 9 (Refeeding) using the following model for each measure of interest: glm(value ~ Geno + TimeOfDay + Phase).

**SOFTWARE USED**
RStudio
bedtools
samtools
Prism

python
CalR (https://calrapp.org/)
hisat2
bowtie2
10XGenomics/Cell Ranger
10XGenomics/subset-bam
Genrich
Macs2
GNU Parallel
R
R packages: rsubread, qvalue, edgeR, limma, plotgardener, bedtoolsr, vcd, bedr
subread/1.5.3
MEME suite SEA
Seurat (Bioconductor)
Metascape (https://metascape.org/)
ClusterCompare (Bioconductor)
Mosaic (Bioconductor)

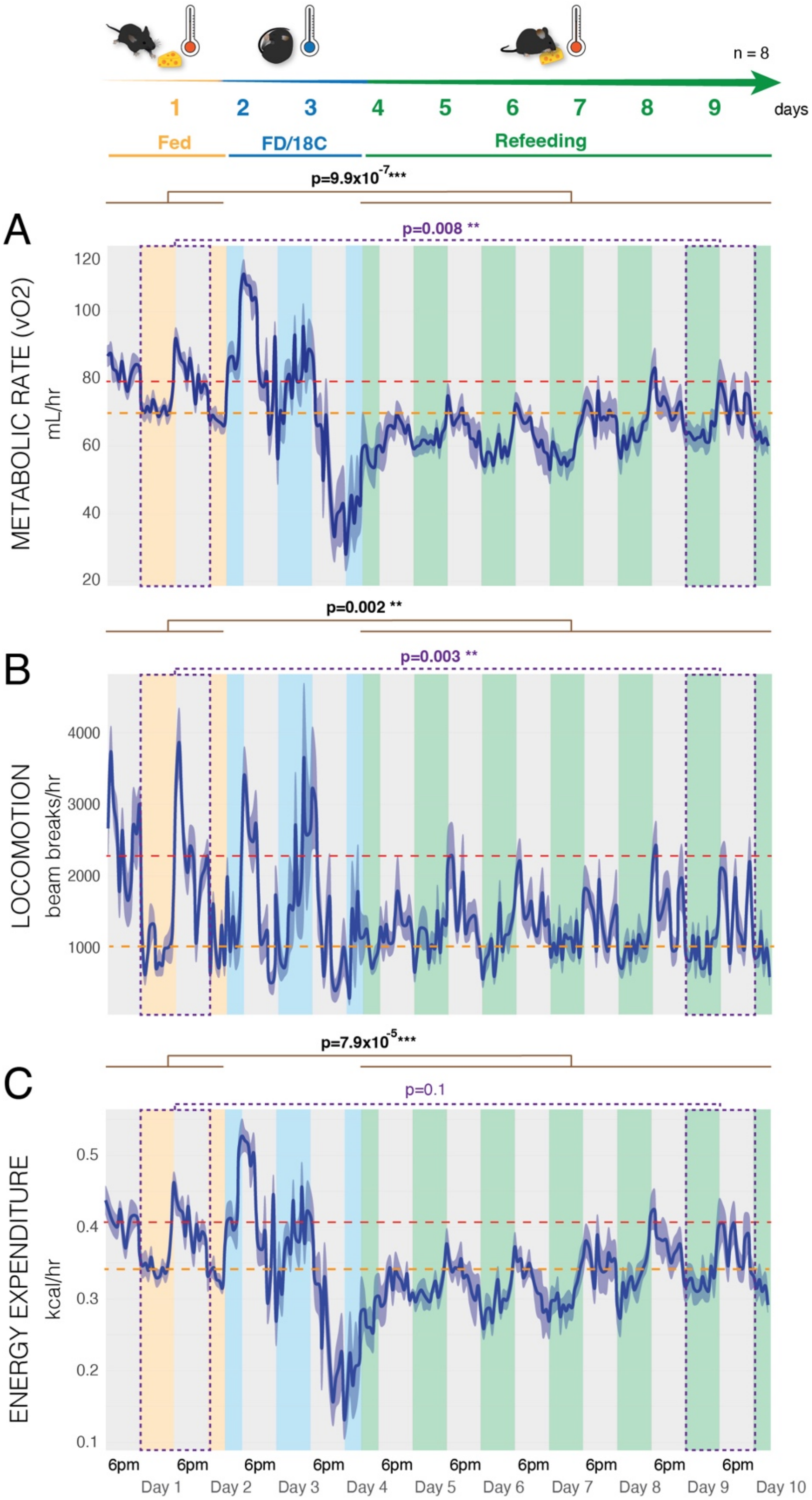
n = 8
1 2 3 4 5 6 7 8 9 days
Fed
FD/18C
Refeeding
p=9.9x10^-7***
A
p=0.008 **
METABOLIC RATE (vO2)
mL/hr
p=0.002 **
B
p=0.003 **
LOCOMOTION
beam breaks/hr
p=7.9x10^-5***
C
p=0.1
ENERGY EXPENDITURE
kcal/hr
6pm
Day 1
Day 2
Day 3
Day 4
Day 5
Day 6
Day 7
Day 8
Day 9
Day 10

**Fig. S1. Persistent metabolic changes in adult mice are observed during the refeeding phase after food deprivation and cold induced torpor.**

**(A-C)** Plots show Metabolic Rate (A), Locomotion (B), Energy Expenditure (C) in adult female mice during Fed (ambient temperature of 24°C) (yellow), food deprived (FD) + cold (ambient temperature of 18°C) (blue), Refeeding periods (green) over 10 days in CLAMS cages. A significant difference between the Fed and Refed states is observed for all three measures (generalized linear model, brown lines), and for the day 1 versus day 9 comparisons for metabolic rate and locomotion (purple). Red dashed line, mean for dark cycle (grey bars). Orange dashed line, mean for light cycle (colored bars). Dark blue line and shading, mean±SEM, n=8.

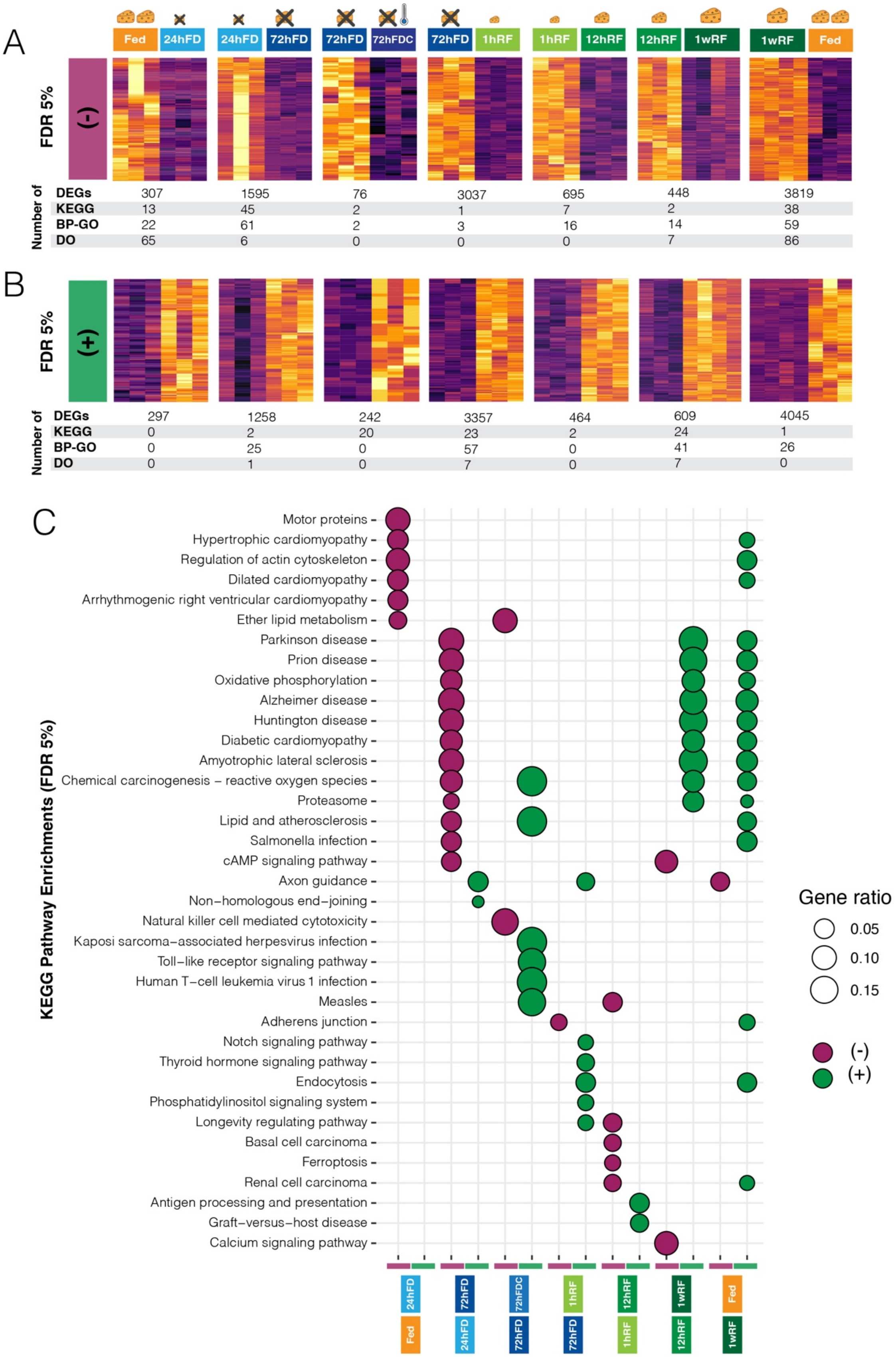
A
Fed
24hFD
24hFD
72hFD
72hFD
72hFDC
72hFD
1hRF
1hRF
12hRF
12hRF
1wRF
1wRF
Fed
FDR 5%
(-)
Number of
DEGs 307 1595 76 3037 695 448 3819
KEGG 13 45 2 1 7 2 38
BP-GO 22 61 2 3 16 14 59
DO 65 6 0 0 0 7 86
B
FDR 5%
(+)
Number of
DEGs 297 1258 242 3357 464 609 4045
KEGG 0 2 20 23 2 24 1
BP-GO 0 25 0 57 0 41 26
DO 0 1 0 7 0 7 0
C
KEGG Pathway Enrichments (FDR 5%)
Motor proteins
Hypertrophic cardiomyopathy
Regulation of actin cytoskeleton
Dilated cardiomyopathy
Arrhythmogenic right ventricular cardiomyopathy
Ether lipid metabolism
Parkinson disease
Prion disease
Oxidative phosphorylation
Alzheimer disease
Huntington disease
Diabetic cardiomyopathy
Amyotrophic lateral sclerosis
Chemical carcinogenesis – reactive oxygen species
Proteasome
Lipid and atherosclerosis
Salmonella infection
cAMP signaling pathway
Axon guidance
Non–homologous end–joining
Natural killer cell mediated cytotoxicity
Kaposi sarcoma–associated herpesvirus infection
Toll–like receptor signaling pathway
Human T–cell leukemia virus 1 infection
Measles
Adherens junction
Notch signaling pathway
Thyroid hormone signaling pathway
Endocytosis
Phosphatidylinositol signaling system
Longevity regulating pathway
Basal cell carcinoma
Ferroptosis
Renal cell carcinoma
Antigen processing and presentation
Graft–versus–host disease
Calcium signaling pathway
24hFD
Fed
72hFD
24hFD
72hFDC
72hFD
1hRF
72hFD
12hRF
1hRF
1wRF
12hRF
Fed
1wRF
Gene ratio
0.05
0.10
0.15
(-)
(+)

**Fig. S2. Different pathways, biological processes, and disease processes are affected at each step of the FFR response in the mouse hypothalamus.**

**(A and B)** Heatmaps show significant differentially expressed genes (DEGs) that decrease (A, purple) or increase (B, green) their expression at each step of the FFR response program (contrasted conditions shown on the top). The table under the heatmaps shows the numbers of significant DEGs (FDR < 5%) and numbers of significantly enriched KEGG pathways, biological processes (BP-GO), and disease ontology terms for each step (FDR < 5%).

**(C)** The plot shows significantly enriched KEGG pathways for increased (green) and decreased (purple) FFR response DEGs. Plotted circles are sized according to the proportion of DEGs found among the genes in the pathway. The results show how different pathways are affected at each step of the FFR response in the hypothalamus.

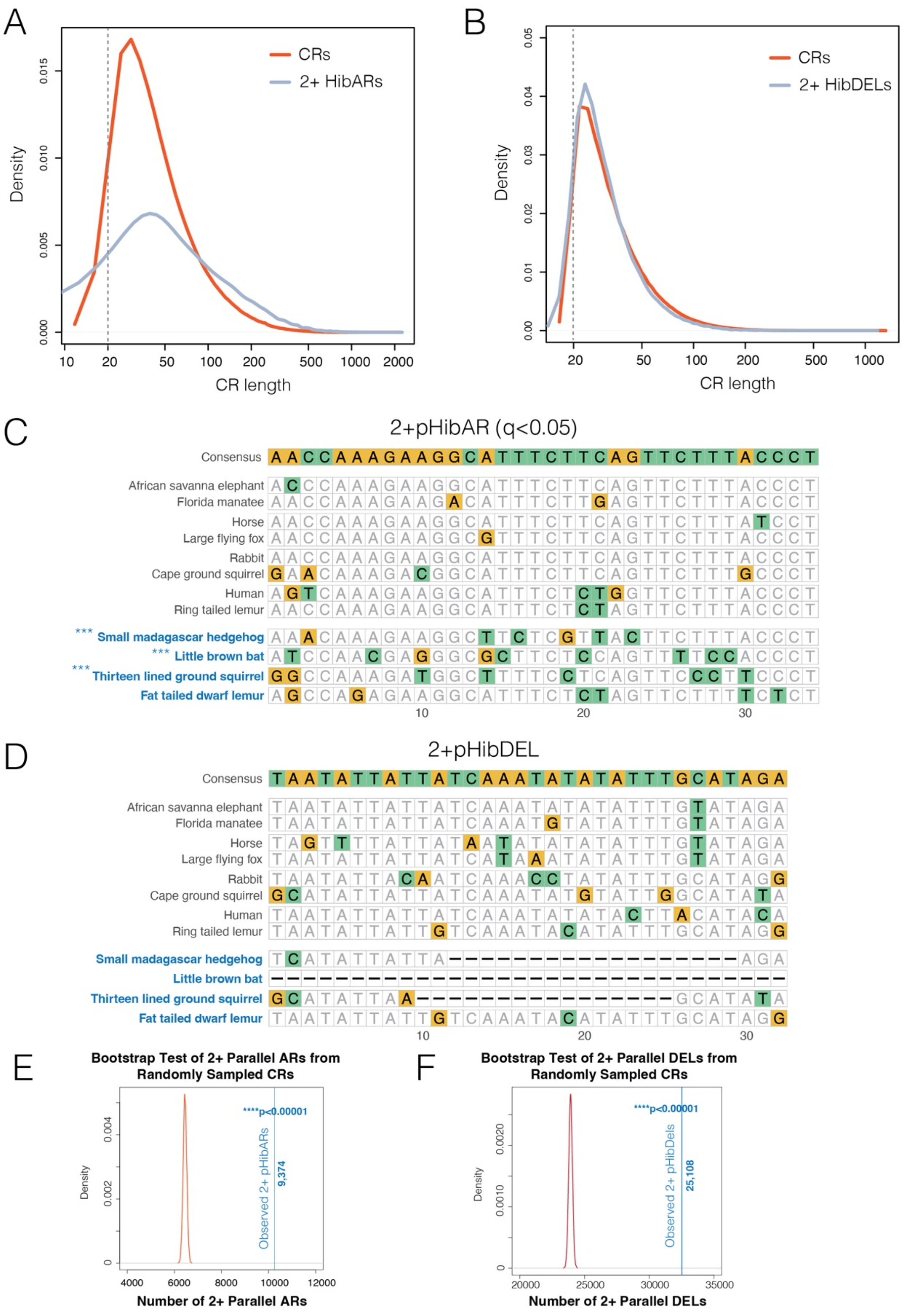

A
Density
CR length
CRs
2+ HibARs
0.000
0.005
0.010
0.015
10
20
50
100
200
500
1000
2000
B
Density
CR length
CRs
2+ HibDELs
0.00
0.01
0.02
0.03
0.04
0.05
20
50
100
200
500
1000
C
2+pHibAR (q<0.05)
Consensus AACCAAAGAAGGCATTTCTTCAGTTCTTTACCCT
African savanna elephant
Florida manatee
Horse
Large flying fox
Rabbit
Cape ground squirrel
Human
Ring tailed lemur
*** Small madagascar hedgehog
*** Little brown bat
*** Thirteen lined ground squirrel
Fat tailed dwarf lemur
10
20
30
D
2+pHibDEL
Consensus TAATATTATTATCAAATATATATTTGCATAGA
African savanna elephant
Florida manatee
Horse
Large flying fox
Rabbit
Cape ground squirrel
Human
Ring tailed lemur
Small madagascar hedgehog
Little brown bat
Thirteen lined ground squirrel
Fat tailed dwarf lemur
10
20
30
E
Bootstrap Test of 2+ Parallel ARs from Randomly Sampled CRs
****p<0.00001
Observed 2+ pHibARs
9,374
Density
0
0.002
0.004
4000
6000
8000
10000
12000
Number of 2+ Parallel ARs
F
Bootstrap Test of 2+ Parallel DELs from Randomly Sampled CRs
****p<0.00001
Observed 2+ pHibDels
25,108
Density
0
0.0010
0.0020
20000
25000
30000
35000
Number of 2+ Parallel DELs

**Fig. S3. Optimized parameters for genome-wide parallel AR and DEL detection uncover convergent sites of accelerated evolution and deletion that exceed chance in hibernators.**
**(A and B)** The density plots show the size distributions (base pairs) of the background conserved regions identified by PhastCons and the size distribution of the CRs identified as significant pHibARs (A) and pHibDELs (B) in 2 or more hibernators (2+ HibARs, blue line). A minimum floor size of 20bp was enforced for the detection of significant ARs (grey line). AR analysis median widths: CRs, 55bp; 2+ pHibARs, 103bp; DEL analysis median widths: CRs, 35bp; 2+ pHibDELs, 32bp.
(**C and D**) The multiple alignments show an example of a pHibAR (C) and pHibDEL (D). The hibernating lineages are indicated in blue type, and the non-hibernator background lineages in black type. The consensus sequence is shown on the top for each example. The results in (C) show conservation of the sequence across the non-hibernating species but significant accelerated evolution in 3 of 4 hibernators, including the small Madagascar hedgehog, little brown bat, and 13-lined ground squirrel. In D, the sequence is conserved across non-hibernators, but 3 hibernating lineages show deleted regions including the Madagascar hedgehog, little brown bat, and 13-lined ground squirrel. ***FDR <0.001 for accelerated evolution in C.
(**E and F**) The density plots show the results of a bootstrap statistical test determining whether the same AR sites (E) and DEL sites (F) were detected in two or more (2+ parallel) hibernators than expected by chance. Random sampling of CRs with replacement shows the distribution of the numbers of 2+ parallel CRs uncovered by chance (red line) compared to the observed number of 2+ parallel ARs (E) or DELs (F) for hibernators (blue line). Significantly more 2+ pHibARs and 2+ pHibDELs are detected from hibernators than expected by chance ($p<0.0001$).

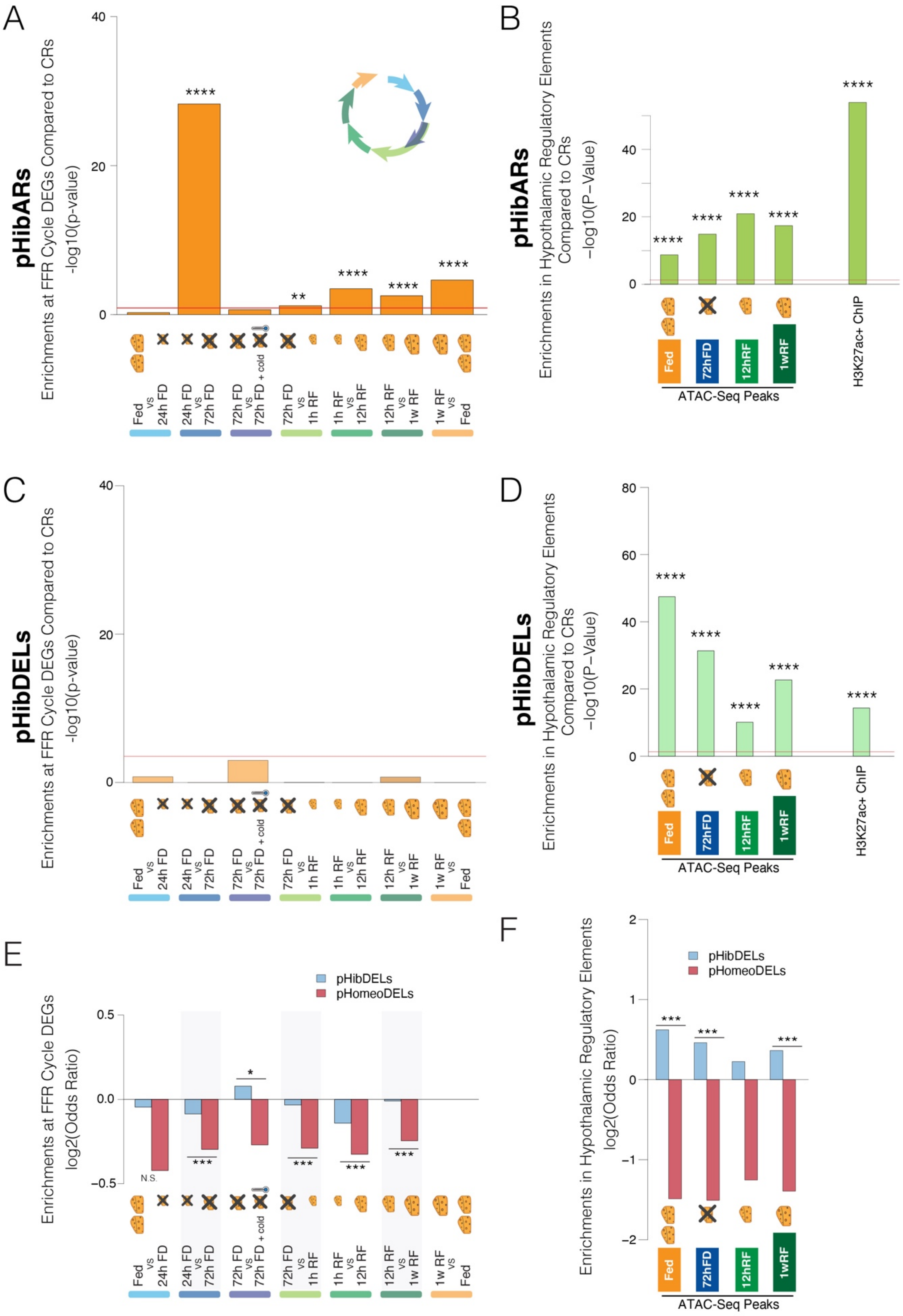
A
pHibARs
Enrichments at FFR Cycle DEGs Compared to CRs
-log10(p-value)
40
20
0
****
**
****
****
****
Fed vs 24h FD
24h FD vs 72h FD
72h FD vs 72h FD + cold
72h FD vs 1h RF
1h RF vs 12h RF
12h RF vs 1w RF
1w RF vs Fed
B
pHibARs
Enrichments in Hypothalamic Regulatory Elements
Compared to CRs
−log10(P−Value)
40
20
0
****
****
****
****
****
Fed
72hFD
12hRF
1wRF
ATAC-Seq Peaks
H3K27ac+ ChIP
C
pHibDELs
Enrichments at FFR Cycle DEGs Compared to CRs
-log10(p-value)
40
20
0
Fed vs 24h FD
24h FD vs 72h FD
72h FD vs 72h FD + cold
72h FD vs 1h RF
1h RF vs 12h RF
12h RF vs 1w RF
1w RF vs Fed
D
pHibDELs
Enrichments in Hypothalamic Regulatory Elements
Compared to CRs
−log10(P−Value)
80
60
40
20
0
****
****
****
****
****
Fed
72hFD
12hRF
1wRF
ATAC-Seq Peaks
H3K27ac+ ChIP
E
Enrichments at FFR Cycle DEGs
log2(Odds Ratio)
pHibDELs
pHomeoDELs
0.5
0.0
−0.5
N.S.
***
*
***
***
***
Fed vs 24h FD
24h FD vs 72h FD
72h FD vs 72h FD + cold
72h FD vs 1h RF
1h RF vs 12h RF
12h RF vs 1w RF
1w RF vs Fed
F
Enrichments in Hypothalamic Regulatory Elements
log2(Odds Ratio)
pHibDELs
pHomeoDELs
2
1
0
−1
−2
***
***
***
Fed
72hFD
12hRF
1wRF
ATAC-Seq Peaks

**Fig. S4. Hibernation-linked genomic changes are disproportionately enriched in regions contacting hypothalamic FFR response DEGs and in *cis*-elements.**

(**A and B**) The barplots show the -log10 of the p-value for enrichment testing of pHibARs versus background CRs (A) in sites forming contacts with FFR response DEGs (A) and in significant FFR response ATAC-Seq peaks and in fed hypothalamus H3K27ac+ ChIP-Seq peaks (B) detected in the hypothalamus of adult mice. Red line shows p=0.05 threshold for significance. FD, food deprived; RF, refed; 12hrs RF, refed for 12hrs after 72 hrs of FD; 1w RF, refed for 1 week after 72hrs of FD; h, hours; w, week; Cold is ambient exposure to 18°C compared to 24°C. Fisher's Exact Test. ****p<0.00001, **p<0.01.

(**C and D**) The barplots show the -log10 of the p-value for enrichment testing of pHibDELs versus background CRs (A) in 10kb windows forming contacts with FFR response DEGs (A) and in significant FFR response ATAC-Seq peaks and H3K27ac+ ChIP-Seq peaks (B) detected in the hypothalamus of adult mice. Red line shows p=0.05 threshold for significance. Fisher's Exact Test. ****p<0.00001, **p<0.01.

(**E**) The barplot shows the odds ratio for pHibDELs (blue) versus control pHomeoDELs (red) forming regulatory contacts with FFR cycle DEGs. Comparisons of pHibDEL versus pHomeoDEL prevalence at each DEG set revealed significantly increased enrichments for pHibDELs over pHomeoDELs at genes involved in the transition from the starvation response (72 hr FD) to the starvation + cold response (72hr FD+cold) (Woolf Test). Additionally, although depleted relative to background CRs, pHibDELs have a significantly greater odds ratio of contacts with FFR cycle DEGs compared to pHomeoDELs for all other FFR cycle steps except the Fed vs 24hr FD contrast. N.S., not significant; *p<0.05, ***p<0.0001

(**F**) The barplot shows the odds ratio of pHibDELs (blue) versus control pHomeoDELs (red) occurring in significant FFR response ATAC-Seq peaks and in fed hypothalamus H3K27ac+ ChIP-Seq peaks compared to background conserved regions (CRs). pHibDELs are significantly enriched compared to pHomeoDELs for Fed, 72hr FD, and 1w RF conditions (Woolf test). ***$p<0.0001$.

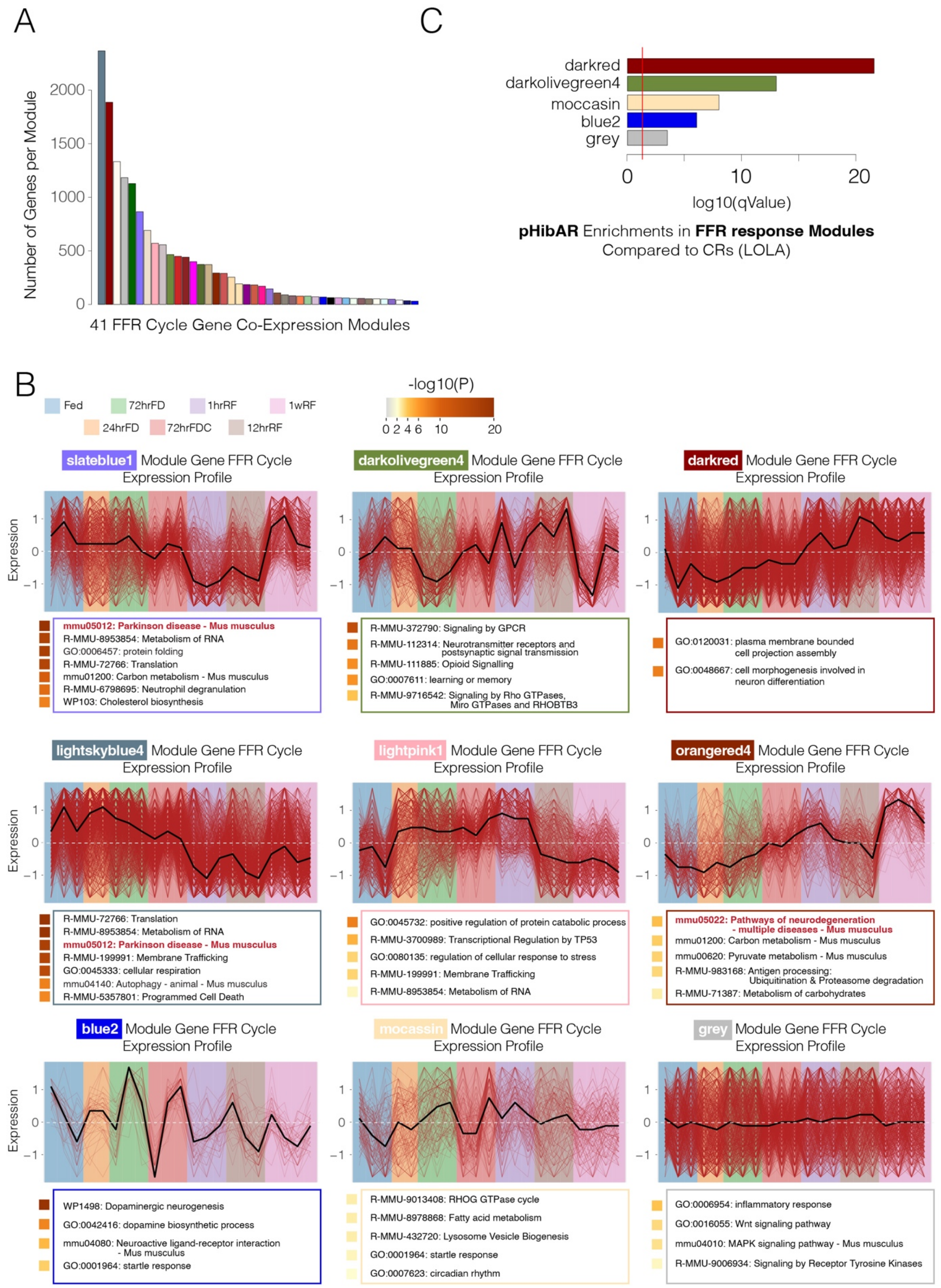
A
Number of Genes per Module
2000
1500
1000
500
0
41 FFR Cycle Gene Co-Expression Modules
C
darkred
darkolivegreen4
moccasin
blue2
grey
0
10
20
log10(qValue)
pHibAR Enrichments in FFR response Modules
Compared to CRs (LOLA)
B
Fed
72hrFD
1hrRF
1wRF
24hrFD
72hrFDC
12hrRF
-log10(P)
0 2 4 6 10 20
slateblue1 Module Gene FFR Cycle Expression Profile
Expression
mmu05012: Parkinson disease - Mus musculus
R-MMU-8953854: Metabolism of RNA
GO:0006457: protein folding
R-MMU-72766: Translation
mmu01200: Carbon metabolism - Mus musculus
R-MMU-6798695: Neutrophil degranulation
WP103: Cholesterol biosynthesis
darkolivegreen4 Module Gene FFR Cycle Expression Profile
R-MMU-372790: Signaling by GPCR
R-MMU-112314: Neurotransmitter receptors and postsynaptic signal transmission
R-MMU-111885: Opioid Signalling
GO:0007611: learning or memory
R-MMU-9716542: Signaling by Rho GTPases, Miro GTPases and RHOBTB3
darkred Module Gene FFR Cycle Expression Profile
GO:0120031: plasma membrane bounded cell projection assembly
GO:0048667: cell morphogenesis involved in neuron differentiation
lightskyblue4 Module Gene FFR Cycle Expression Profile
R-MMU-72766: Translation
R-MMU-8953854: Metabolism of RNA
mmu05012: Parkinson disease - Mus musculus
R-MMU-199991: Membrane Trafficking
GO:0045333: cellular respiration
mmu04140: Autophagy - animal - Mus musculus
R-MMU-5357801: Programmed Cell Death
lightpink1 Module Gene FFR Cycle Expression Profile
GO:0045732: positive regulation of protein catabolic process
R-MMU-3700989: Transcriptional Regulation by TP53
GO:0080135: regulation of cellular response to stress
R-MMU-199991: Membrane Trafficking
R-MMU-8953854: Metabolism of RNA
orangered4 Module Gene FFR Cycle Expression Profile
mmu05022: Pathways of neurodegeneration - multiple diseases - Mus musculus
mmu01200: Carbon metabolism - Mus musculus
mmu00620: Pyruvate metabolism - Mus musculus
R-MMU-983168: Antigen processing: Ubiquitination & Proteasome degradation
R-MMU-71387: Metabolism of carbohydrates
blue2 Module Gene FFR Cycle Expression Profile
WP1498: Dopaminergic neurogenesis
GO:0042416: dopamine biosynthetic process
mmu04080: Neuroactive ligand-receptor interaction - Mus musculus
GO:0001964: startle response
mocassin Module Gene FFR Cycle Expression Profile
R-MMU-9013408: RHOG GTPase cycle
R-MMU-8978868: Fatty acid metabolism
R-MMU-432720: Lysosome Vesicle Biogenesis
GO:0001964: startle response
GO:0007623: circadian rhythm
grey Module Gene FFR Cycle Expression Profile
GO:0006954: inflammatory response
GO:0016055: Wnt signaling pathway
mmu04010: MAPK signaling pathway - Mus musculus
R-MMU-9006934: Signaling by Receptor Tyrosine Kinases

**Fig. S5. FFR response gene co-expression modules uncovered in the adult mouse hypothalamus.**

**(A)** The barplot shows the number of genes in each of the 41 FFR response gene co-expression modules uncovered by RNA-Seq in adult mouse hypothalamus (see Figure 1C).

**(B)** The plots show the expression dynamics of genes in a subset of the different FFR response gene co-expression modules. The back line shows the mean for all genes in the module. Below each module are select top gene ontology, reactome, and KEGG pathway enrichments found for the genes in the modules compared to genes expressed in the hypothalamus (FDR < 5%). The results show how different fed, food deprived (FD), FD+cold (FDC), and refed (RF) states affect each gene module and the processes involved. H, hr; w, week.

**(C)** The barplot shows the -log10 of the q-value for pHibAR and pHomeoAR enrichments at genes in each of the 41 co-expression modules compared to background CRs (LOLA R package). Only the modules significantly enriched are shown and the results show that 5 modules are disproportionately enriched for pHibARs. pARs were assigned to genes based on PLAC-Seq data. Red line shows p=0.05.

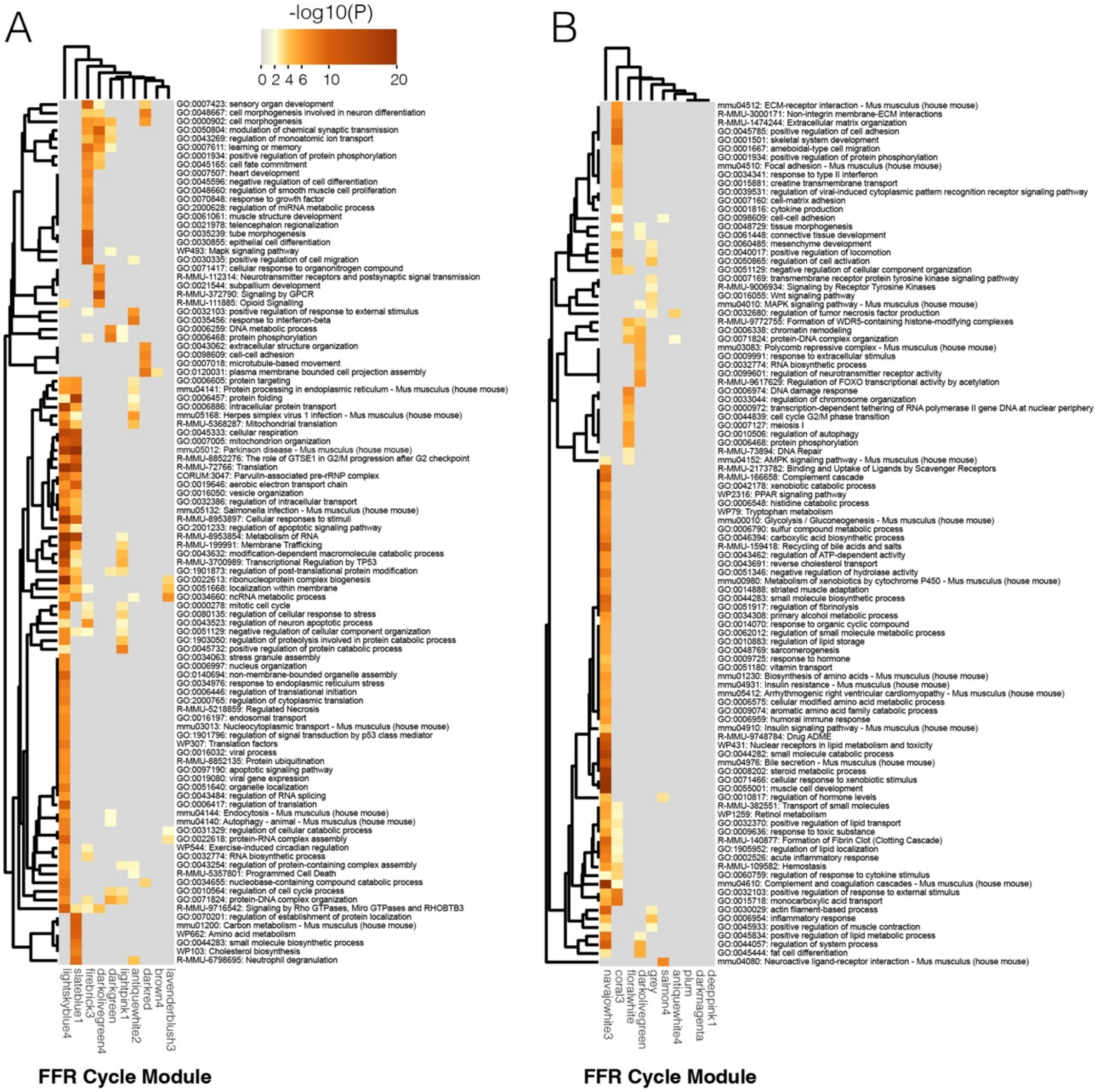
A
B
-log10(P)
0 2 4 6 10 20
FFR Cycle Module
FFR Cycle Module

**Fig. S6. Metascape gene set analysis results for FFR co-expression modules 1-20.**

The heatmaps show the results of a comparative gene set enrichment analysis for FFR response gene co-expression modules 1-10 (**A**) and 11-20 (**B**) using Metascape. The name for each module is shown on the x-axis. The top 100 significantly enriched pathways and biological processes are shown for each gene co-expression module.

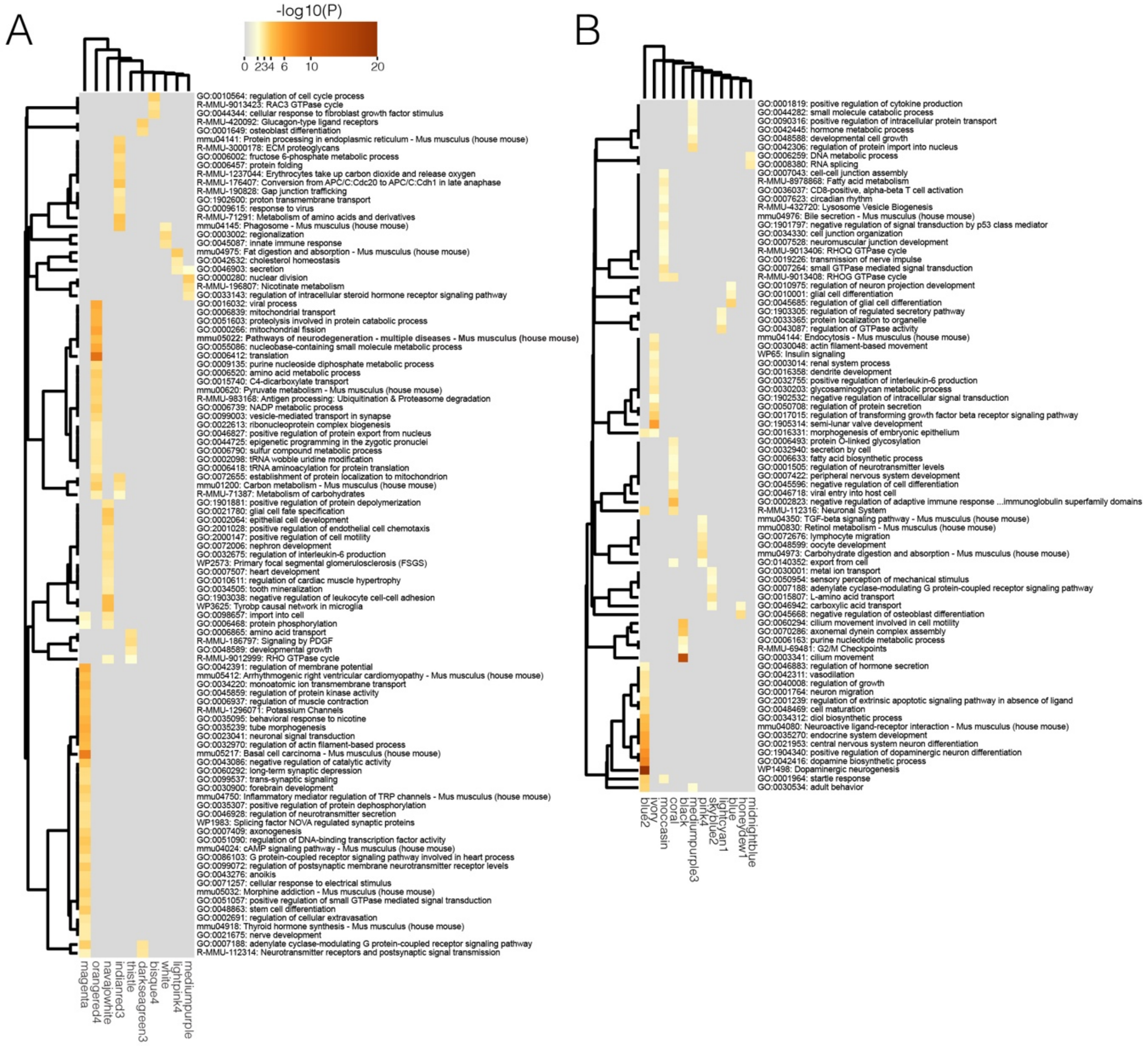
A
B
-log10(P)
0 2 3 4 6 10 20
magenta
orangered4
navajowhite
indianred3
thistle
darkseagreen3
bisque4
white
lightpink4
mediumpurple
blue2
ivory
moccasin
coral
black
mediumpurple3
pink4
skyblue2
lightcyan1
blue
honeydew1
midnightblue

**Fig. S7. Metascape gene set analysis results for FFR co-expression modules 21-41.**

The heatmaps show the results of a comparative gene set enrichment analysis for FFR response gene co-expression modules 21-30 (**A**) and 31-41 (**B**) using Metascape. The name for each module is shown on the x-axis. The top 100 significantly enriched pathways and biological processes are shown for each gene co-expression module.

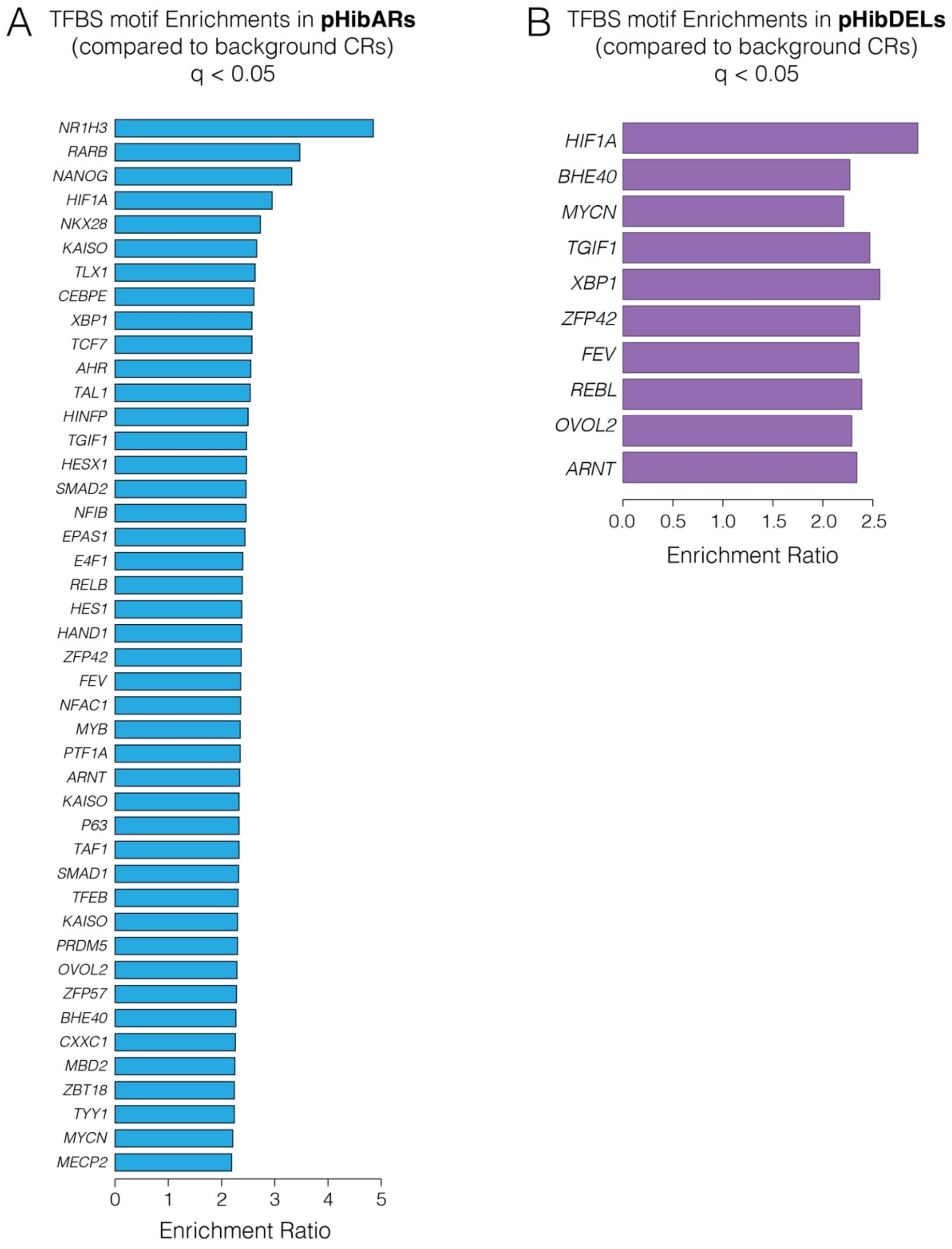
A
TFBS motif Enrichments in **pHibARs**
(compared to background CRs)
q < 0.05
NR1H3
RARB
NANOG
HIF1A
NKX28
KAISO
TLX1
CEBPE
XBP1
TCF7
AHR
TAL1
HINFP
TGIF1
HESX1
SMAD2
NFIB
EPAS1
E4F1
RELB
HES1
HAND1
ZFP42
FEV
NFAC1
MYB
PTF1A
ARNT
KAISO
P63
TAF1
SMAD1
TFEB
KAISO
PRDM5
OVOL2
ZFP57
BHE40
CXXC1
MBD2
ZBT18
TYY1
MYCN
MECP2
0
1
2
3
4
5
Enrichment Ratio
B
TFBS motif Enrichments in **pHibDELs**
(compared to background CRs)
q < 0.05
HIF1A
BHE40
MYCN
TGIF1
XBP1
ZFP42
FEV
REBL
OVOL2
ARNT
0.0
0.5
1.0
1.5
2.0
2.5
Enrichment Ratio

**Fig. S9. Human orthologs for hibernation-linked genomic changes enrich motifs that point to upstream regulatory pathways.**

Barplots show the enrichment ratios for transcription factor binding site motifs (TFBS) in human orthologous sequences for pHibARs (**A**) and pHibDELs (**B**) compared to background CRs. All shown enrichments are statistically significant (FDR < 5%). The results point to upstream regulatory proteins acting on pHibAR and pHibDEL sites in the human genome.

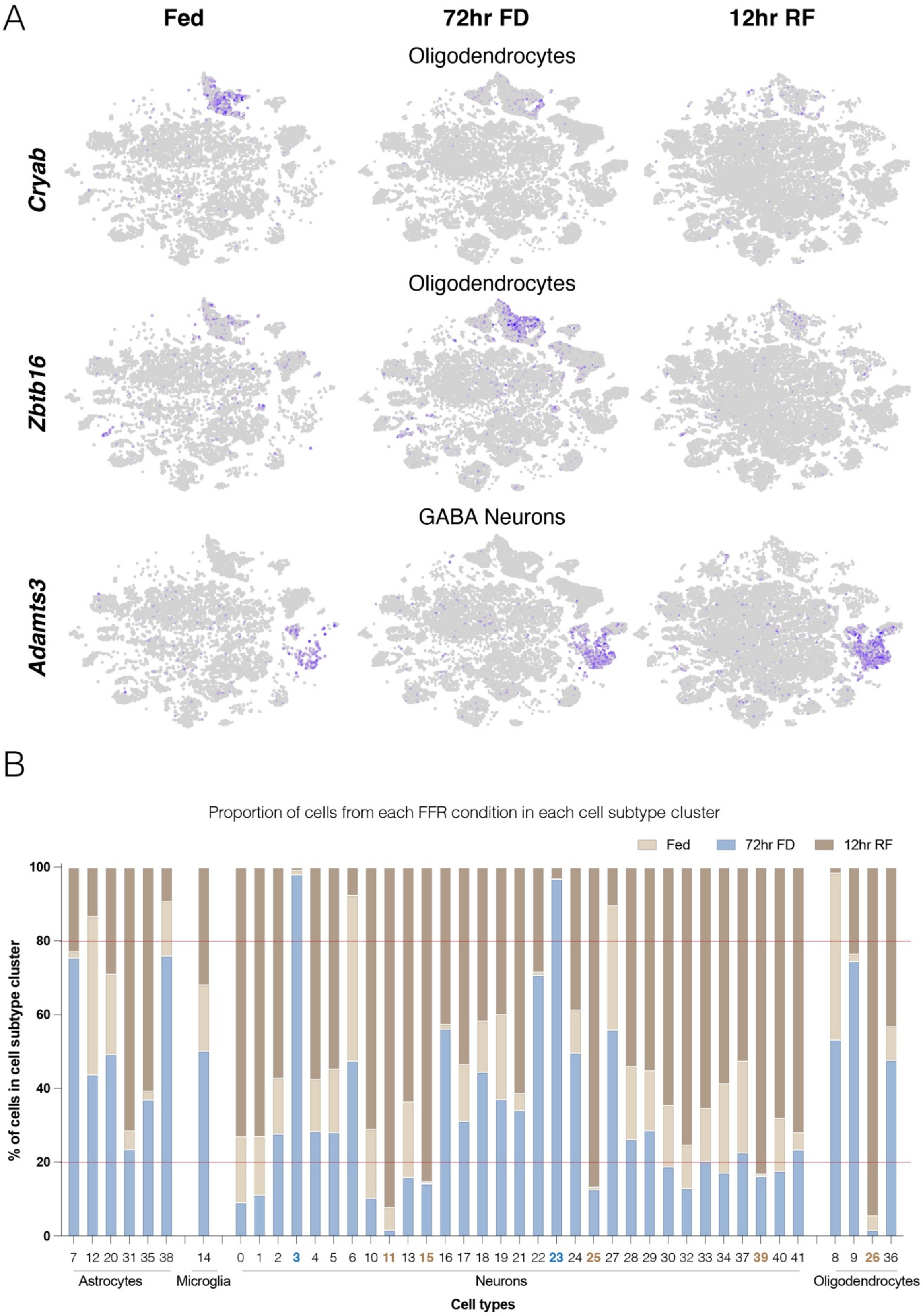
A
Fed
72hr FD
12hr RF
Oligodendrocytes
Cryab
Oligodendrocytes
Zbtb16
GABA Neurons
Adamts3
B
Proportion of cells from each FFR condition in each cell subtype cluster
Fed
72hr FD
12hr RF
100
80
60
40
20
0
% of cells in cell subtype cluster
7 12 20 31 35 38
14
0 1 2 3 4 5 6 10 11 13 15 16 17 18 19 21 22 23 24 25 27 28 29 30 32 33 34 37 39 40 41
8 9 26 36
Astrocytes
Microglia
Neurons
Oligodendrocytes
Cell types

**Fig. S10. Single cell RNA-Seq identifies FFR changes to gene expression at the cellular level in the hypothalamus.**

**(A)** The tSNE plots show examples of genes with expression changes in the hypothalamus at the cellular level in Fed, 72hr FD, and 12hr RF conditions. The gene, *Cryab,* shows elevated expression in Fed, but reduced expression in 72hr FD and 12 hr RF oligodendrocytes, while *Zbtb16* shows elevated expression in the 72 hr FD condition. The gene, *Adamts3,* is an example of a gene that shows elevated expression in the 12hr RF condition in GABA neurons.

(**B**) The barplot shows the percentage of cells found in each FFR condition (Fed, 72hr FD, 12hr RF) for each of 41 different cell subtypes. The results reveal molecular cell subtypes that predominate in particular FFR states, such as the 72hr FD (blue) or 12hr RF (dark brown) states.

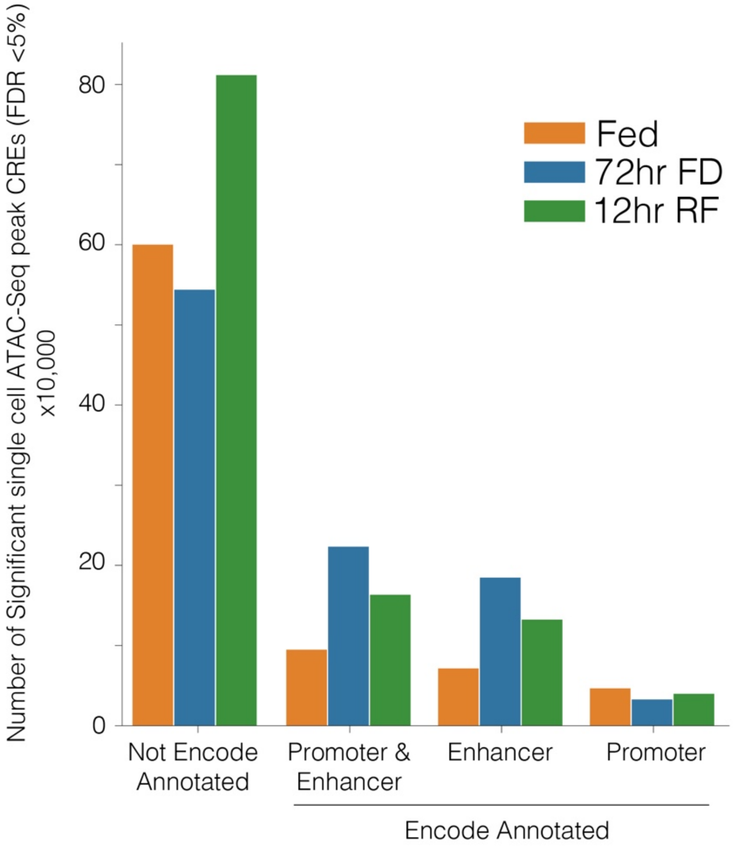
Number of Significant single cell ATAC-Seq peak CREs (FDR <5%) x10,000
80
60
40
20
0
Fed
72hr FD
12hr RF
Not Encode Annotated
Promoter & Enhancer
Enhancer
Promoter
Encode Annotated

**Fig. S11. Comparison of hypothalamic CREs uncovered by single cell ATAC-Seq for FFR states to ENCODE annotated CREs.**

The barplot shows the number of significant single cell ATAC-Seq peaks (FDR < 5%) found from all cell types in the mouse hypothalamus according to whether they overlap ENCODE annotated enhancers, promoters, or sites annotated as both enhancers and promoters. Many sites do not overlap known ENCODE CREs, especially those uncovered in the 12hr RF condition, suggesting new putative active CREs.

### SUPPLEMENTAL TABLES

**Table S1. FFR Response Significant DEGs.** Excel sheets of significant differentially expressed genes in the adult mouse hypothalamus for each FFR step (FDR 5%; EdgeR package).

**Table S2. FFR Response DEG Gene Set Analysis.** Gene set enrichments for FFR response DEGs using the ClusterCompare R package against the background of expressed genes in the hypothalamus (FDR 5%). KEGG (tab1), Reactome (tab2), Gene Ontology (tab3), and Disease Ontology (tab4) enrichments are shown. The FFR response condition contrasts are indicated in the Contrast column, with the direction of the expression change for the second condition indicated (up versus down change in expression level).

**Table S3. FFR Response Consensus ATAC-Seq peak sites.** Bedfiles of consensus ATAC-Seq peaks for FFR responses in mm10 coordinates (FDR <5% in 4 of 8 replicates), including fed (tab1), 72hr FD (tab2), 12hr refed (tab3), and 1 week refed (tab4).

**Table S4. Genomic coordinates (mm10) and statistical results for significant ARs for each hibernator lineage in background conserved regions.**

**Table S5. Significant pHibARs, pHibDels, pHomeoARs, and pHomeoDELs in hg38 coordinates.**

**Table S6. H3K27ac+ PLAC-Seq results connecting *cis*-elements to gene promoters in the mouse hypothalamus (10kb cis-bins).**

**Table S7. H3K27ac+ Hypothalamus PLAC-Seq assignments of pHibARs and pHibDELs to contacted gene promoters.**

**Table S8. FFR Response Gene Co-Expression Module Hub Genes and pHibAR-Hub genes and Reactome Pathway Enrichment Results.**

**Table S9. Cell type and subtype cluster marker genes for cell populations identified in the adult female hypothalamus using scRNA-Seq.**

**Table S10. pHibAR-Hub Genes that are also significant marker genes for hypothalamic cell subtype clusters.**

**Data S1. Fed Hibernation-Linked Hypothalamus Cellular *Cis*-Regulatory Programs.** The ZIP file directory contains BED files and tables of scATAC-Seq results showing CRE activity in 8 different hypothalamic cell types in Fed mice, overlaps with pHibARs and/or pHibDELs, and associated genes with PLAC-Seq detected regulatory contacts.

**Data S2. 72-hr Food Deprived Hibernation-Linked Hypothalamus Cellular *Cis*-Regulatory Programs.** The ZIP file directory contains BED files and tables of scATAC-Seq results showing

CRE activity in 8 different hypothalamic cell types in 72-hr food deprived mice, overlaps with pHibARs and/or pHibDELs, and associated genes with PLAC-Seq detected regulatory contacts.

**Data S3. 12-hr Refeeding Hibernation-Linked Hypothalamus Cellular *Cis*-Regulatory Programs.** The ZIP file directory contains BED files and tables of scATAC-Seq results showing CRE activity in 8 different hypothalamic cell types in 12-hr refed mice following 72-hr food deprivation, overlaps with pHibARs and/or pHibDELs, and associated genes with PLAC-Seq detected regulatory contacts.
